# Matrix architecture and mechanics regulate myofibril organization, costamere assembly, and contractility of engineered myocardial microtissues

**DOI:** 10.1101/2023.10.20.563346

**Authors:** Samuel J. DePalma, Javiera Jillberto, Austin E. Stis, Darcy D. Huang, Jason Lo, Christopher D. Davidson, Aamilah Chowdhury, Maggie E. Jewett, Hiba Kobeissi, Christopher S. Chen, Emma Lejeune, Adam S. Helms, David A. Nordsletten, Brendon M. Baker

**Affiliations:** Department of Biomedical Engineering, University of Michigan, Ann Arbor, MI 48109; Department of Mechanical Engineering, Boston University, Boston, MA 02215; Division of Cardiovascular Medicine, University of Michigan, Ann Arbor, MI 48109; Department of Cardiac Surgery, University of Michigan, Ann Arbor, MI 48109; Department of Biomedical Engineering, School of Imaging Sciences and Biomedical Engineering, King’s College London, King’s Health Partners, London SE1 7EH, United Kingdom; Department of Biomedical Engineering, Boston University, Boston, MA 02215, USA; Wyss Institute for Biologically Inspired Engineering, Harvard University, Boston, MA 02115, USA; Department of Chemical Engineering, University of Michigan, Ann Arbor, MI 48109

**Keywords:** biomaterials, electrospinning, cardiac tissue engineering, cardiomyocytes, induced pluripotent stem cells, extracellular matrix, microenvironment, mechanosensing, matrix stiffness

## Abstract

The mechanical function of the myocardium is defined by cardiomyocyte contractility and the biomechanics of the extracellular matrix (ECM). Understanding this relationship remains an important unmet challenge due to limitations in existing approaches for engineering myocardial tissue. Here, we established arrays of cardiac microtissues with tunable mechanics and architecture by integrating ECM-mimetic synthetic, fiber matrices and induced pluripotent stem cell-derived cardiomyocytes (iPSC-CMs), enabling real-time contractility readouts, in-depth structural assessment, and tissue-specific computational modeling. We find that the stiffness and alignment of matrix fibers distinctly affect the structural development and contractile function of pure iPSC-CM tissues. Further examination into the impact of fibrous matrix stiffness enabled by computational models and quantitative immunofluorescence implicates cell-ECM interactions in myofibril assembly and notably costamere assembly, which correlates with improved contractile function of tissues. These results highlight how iPSC-CM tissue models with controllable architecture and mechanics can inform the design of translatable regenerative cardiac therapies.

## INTRODUCTION

Heart disease remains the leading cause of death worldwide^1^. Despite recent advances in treatment, existing therapies for treating heart disease fail to restore normal function of the heart following chronic or acute injury, due in part to the limited regenerative potential of the myocardium^2,3^. Thus, there is a critical need for regenerative or tissue-replacement therapies that restore normal cardiac architecture and mechanical function. In recent years, advances in induced pluripotent stem cell (iPSC) technologies have made the creation of engineered heart tissues (EHTs) feasible for use as regenerative therapies, *in vitro* models to study cardiac regeneration, or screening platforms to test the effectiveness and/or toxicity of new therapeutics^4–6^.

Among the many techniques explored to generate mature iPSC-derived cardiomyocyte (iPSC-CM) tissues, significant efforts have focused on developing scaffolds that recapitulate physiologic tissue organization to improve overall tissue function and potentially maturity^7–9^. The mechanical function of the myocardium is dictated by contractile CMs and the surrounding fibrous extracellular matrix (ECM) that organizes and supports CMs^10–12^. Individual layers of muscle tissue throughout the myocardium are highly anisotropic, driving coordinated uniaxial contractions^13^. These muscle fibers and their accompanying ECM twist transmurally, thus generating the torsional contractile behavior critical to proper systolic function of the ventricle^14^. As such, scaffolds that recapitulate biochemical and mechanical features of native cardiac ECM and direct cellular orientation hold promise for improving the function and maturation of cardiac tissue constructs^15–20^. Scaffolding is often integrated into EHTs by combining naturally derived biomaterials materials such as purified collagen or fibrin with iPSC-CMs^8,20–24^. However, these materials provide limited control over mechanical properties which is critical to providing insight into how iPSC-CMs sense and respond to matrix stiffness. Moreover, the addition of admixed stromal cells is often required to drive proper tissue assembly in these systems, precluding the direct study of CM behavior. Prior studies have explored the use of polymeric hydrogels providing improved mechanical control or electrospun polymeric scaffolds with fibrous topography that better recapitulates the native ECM; however, these materials either lack fibrous topography or sufficient tunability, respectively, both of which are needed for identifying critical mechanobiological mechanisms that driver cardiac tissue assembly or maturation^16,17,19,25–27^. Thus, EHTs formed from iPSC-CMs and highly tunable, fibrous matrix providing orthogonal control over architecture and mechanics could provide new and important insights into how iPSC-CMs respond to physical microenvironment inputs. Furthermore, a deeper understanding of how CMs interact with their physical microenvironment could in turn establish key design attributes of scaffolds that support stem cell-derived cardiac tissue formation and maturation.

The interactions between cardiomyocytes and their surrounding native ECM or a biomaterial scaffold are regulated by cellular mechanosensing and ultimately the transduction of mechanical forces into cell signaling cascades. CM mechanosensing has been shown to be critical in cardiac development, disease progression, and the assembly of *in vitro* engineered heart tissues, highlighting the necessity for the informed design of scaffolds used to engineer mature iPSC-CM tissues^17,28–32^. As CMs have extremely dynamic mechanical functions, they use multiple mechanosensing mechanisms to sense and respond to changes in their mechanical environment^28^. Genetic variants that impact the function of mechanosensing proteins can render cardiomyocytes abnormally susceptible to mechanical stresses, leading to ventricular hypertrophy or dilation and ultimately heart failure^33–35^. In particular, the mechanosensitive protein vinculin is known to play a critical role at both costameres and intercalated discs during both cardiac development and disease^30,36–40^. Genetic variants of vinculin contribute to cardiomyopathies^34^ and cardiomyocyte-specific knock-out of vinculin leads to early cardiac failure or dilated cardiomyopathies^39,40^. Conversely, overexpression of vinculin in drosophila hearts leads to myocardial remodeling and improved cardiac function in aging hearts, resulting in extended organismal lifespan^31^. Moreover, the mechanosensitivity of vinculin in CMs has been described both *in vitro* and *in vivo,* suggesting that vinculin localization to costameres and intercalated discs regulates myofibril maturation and is influenced by mechanical signals^26,30,38,41,42^. Despite these advances in our knowledge of vinculin’s role in CM mechanosensing, it is unknown how fibrous matrix mechanics impact vinculin’s localization to cell-cell or cell-ECM adhesions and the resulting implications on EHT function and maturation.

To study how specific and tissue-relevant structural and mechanical cues impact engineered cardiac tissue assembly, maturation, and function, we established a novel biomaterials-based platform for creating arrays of cardiac microtissues composed of tunable, synthetic, fibrous ECM and purified iPSC-CMs. Through carefully controlled studies varying ECM organization and mechanics, we found that microtissues formed on soft (< 1 kPa), aligned fibrous matrices tethered between soft elastic posts demonstrate improved CM adhesion, organization, and contractile function. Tissue-specific computational modeling revealed that altered cellular mechanical behavior, in conjunction with the passive mechanics of the tissue, drive the observed changes in tissue contractility. Associated with these effects, we found that vinculin localization to costameres during tissue formation was dependent on fibrous matrix stiffness. Moreover, robust vinculin localization to costameres was strongly associated with tissue maturation and increases in contractile function. These findings highlight the importance of highly controlled bioengineered platforms for studying CM mechanosensing and provide several key insights that inform the design of biomaterial scaffolds for engineered cardiac tissue replacement therapies.

## RESULTS

### Development and characterization of mechanically tunable engineered heart tissue platform

As the myocardial microenvironment plays a vital role in both cardiac development and disease progression, constructing EHTs with biomaterials that recapitulate relevant architecture and mechanics of the fibrous cardiac ECM is critical for studying these processes *in vitro*^9^. We previously developed matrices composed of synthetic dextran vinyl sulfone (DVS) polymeric fibers that possess comparable geometry to perimysial collagen fibers, which are ∼1 μm in diameter and surround CM bundles to confer tissue mechanical anisotropy and enable mechanical function of cardiac tissue ^10–12,15^. Using this biomaterial platform, we showed that matrix fiber alignment is critical to driving proper tissue organization and calcium handling dynamics^15^. Additionally, we found that these fibrous scaffolds facilitate long-term culture of iPSC-CMs (>28 days) enabled by robust cell-matrix interactions^15^. However, this culture platform did not allow for tissue fractional shortening, thereby preventing the assessment of contractile function of formed EHTs. Predicated on this previous work, here we advanced this model by generating a platform enabling fractional shortening and orthogonal mechanical tunability to explore the impact of an expanded array of architectural and mechanical cues on iPSC-CM function.

To examine how variations in the architecture and mechanics of the cardiac microenvironment influence iPSC-CM tissue formation, we established a novel approach to generating arrays of cardiac microtissues (termed fibroTUGs or fibrous tissue μ-gauges) composed of tunable electrospun, synthetic fiber matrices suspended between two elastomeric posts seeded with pure populations of iPSC-CMs using a microfabrication-based cell patterning strategy. Microfabricated PDMS post arrays consisting of 98 pairs of rectangular posts were fabricated with standard soft lithography techniques (**Fig. 1a,b**). Based on previous literature^22,43^ and as confirmed via custom mechanical characterization methods (**Fig. S1a-c**), post heights were defined to generate soft (0.41 N/m) and stiff (1.2 N/m) mechanical boundary conditions (**Fig. 1a,h**). Subsequently, electrospun DVS fibers were deposited upon post arrays affixed to a collecting mandrel rotating at various speeds to control fiber alignment, as previously described^15^ (**Fig. 1a,f,g**). Next, fiber matrices spanning two posts were stabilized via photoinitiated free radical crosslinking by exposing substrates to UV light through a photopatterning mask in the presence of lithium phenyl-2,4,6-trimethylbenzoylphosphinate (LAP) photoinitiator. Upon hydration, uncrosslinked fibers were dissolved, resulting in fibrous matrices spanning only between pairs of posts. Stabilized fibrous matrices spanning posts could then be stiffened further via exposure to UV light in the presence of LAP to define a final matrix stiffness^15,44^. Crosslinking parameters were identified to generate matrices with stiffnesses corresponding to developing (0.1 mg/mL LAP; 0.68 kPa), adult (1.0 mg/mL LAP; 10.1 kPa), or diseased (5.0 mg/mL LAP; 17.4 kPa) myocardium, as characterized by microindentation measurements (**Fig. 1i, S1d-f**)^17,45,46^. To generate pure cardiomyocyte tissues without the requirement of admixed stromal cells, purified cultures of iPSC-CMs were seeded upon photopatterned matrices using a physically registered, microfabricated seeding mask that funneled iPSC-CMs to the suspended matrices, limiting seeded cells from settling and adhering to the glass surface below suspended fiber matrices (**Fig. 1a**). The resulting tissues contained ∼100 iPSC-CMs regardless of the pre-defined mechanics of the posts or matrices (**Fig, 1j**).

**Figure 1:**
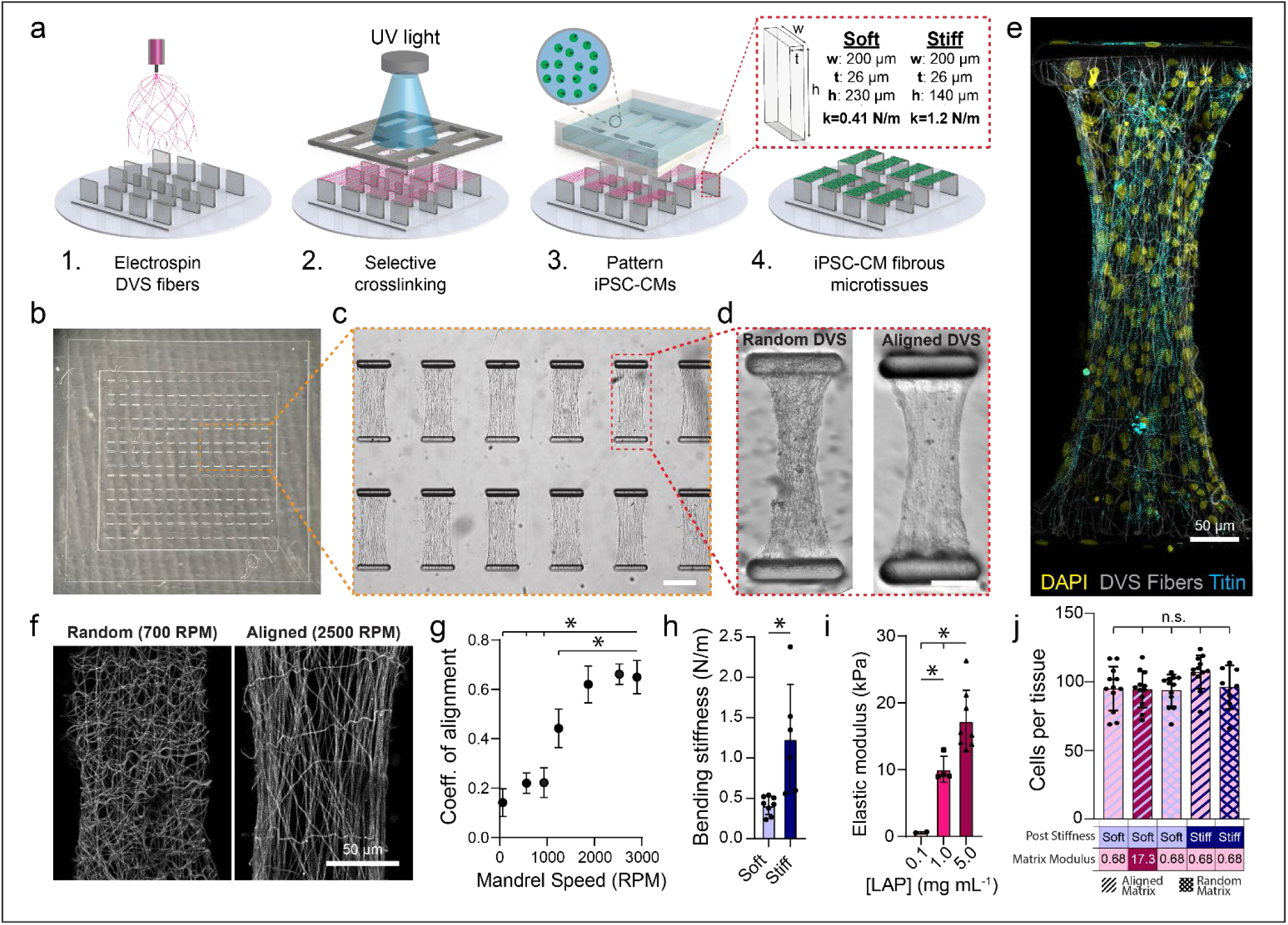
Fabrication of pure iPSC-CM microtissues with high mechanical tunability. (a) Schematic of fibroTUG fabrication and seeding. (b) Full array of microfabricated PDMS posts before DVS fiber electrospinning. (c) Brightfield image of photopatterned suspended matrices (scale bar: 200 μm). (d) Representative brightfield images of pure populations of iPSC-CMs seeded on random and aligned DVS matrices 7 days after seeding (scale bar: 100 μm). (e) Confocal fluorescent image of fibroTUG tissue formed on 0.68 kPa, aligned matrices suspended between soft posts. (f) Confocal fluorescent images of random and aligned fiber matrices functionalized with methacrylated rhodamine. (g) Rotation speed of the collection mandrel during fiber electrospinning was varied to define fiber alignment (n ≥ 5 matrices). (h) Post height was varied to define post bending stiffness. (i) LAP photoinitiator concentration was tuned to generate matrices of physiologically relevant stiffnesses. (j) Tissue seeding was unaffected by the mechanical inputs, as quantified by the number of cells that compose each tissue 7 days after seeding. All data presented as mean ± std; * p < 0.05.

While matrix stiffness has previously been extensively studied using polymeric hydrogel or elastomer surfaces^17,18,26,46,47^, little is known about how the stiffness of assemblies of fibers influences iPSC-CM tissue formation and function. The discrete nature of fibrous matrices engenders distinct behavior compared to elastic, continuum-like materials^48^. Although here we provide measurements of bulk modulus to demonstrate mechanical tunability, these values cannot be directly extrapolated to continuum-like materials. Of note, these matrix fibers are individually and at the nano-scale are quite stiff (∼100’s of MPa) despite overall soft bulk stiffness, in contrast to hydrogel or elastomer materials where mechanics are uniform across multiple length scales. Furthermore, because our fibroTUG platform is predicated upon predefined synthetic matrices that facilitate robust integrin engagement to guide assembly of myocardial syncytia that are structurally similar to stratified muscle layers in the myocardium, we were able to generate and monitor functional myocardial tissues composed of pure iPSC-CMs, highlighting the utility of this system in studying the impact of mechanical cues on cell- and tissue-scale function over long-term culture (up to 2 weeks) (**Fig. S2**)^7^.

### Mechanical inputs impact tissue mechanical function, organization, and maturation

After thorough mechanical characterization of our fibroTUG platform, we examined how altering key architectural and mechanical inputs – specifically matrix alignment, matrix stiffness, and post stiffness – affected iPSC-CM tissue assembly, organization, and function (**Fig. 2**). To assess resulting tissue function, contractility of tissues after 7 days of culture was quantified by measuring post deflections from time-lapse imaging of contracting tissues (**Supp. Videos 1-4**). Myofibril organization and density were quantified as previously described to assess the impact of physical microenvironmental cues on tissue organization^15,49^.

**Figure 2:**
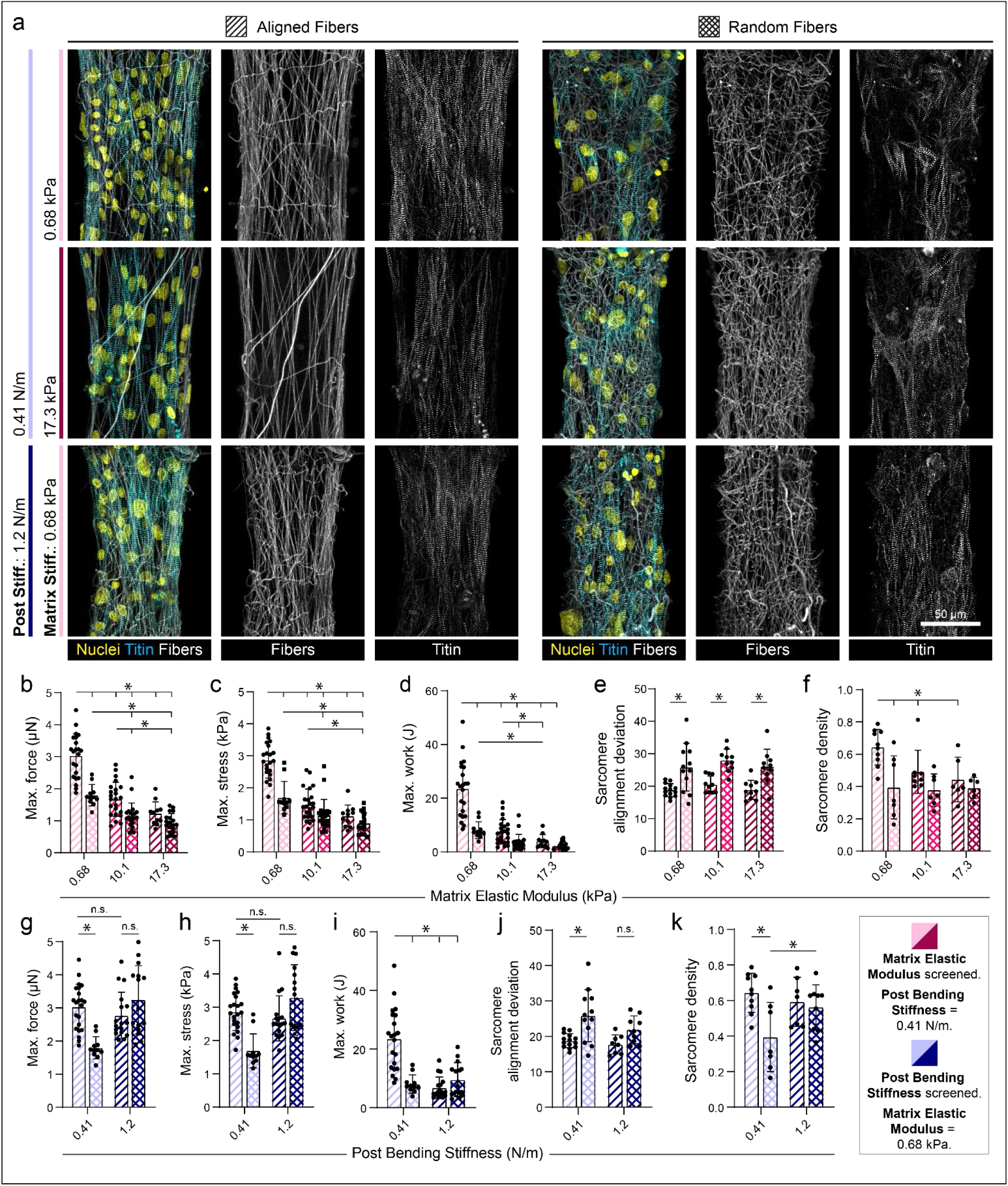
Fibrous matrix alignment and stiffness influences iPSC-CM tissue assembly and force generation. (a) Confocal fluorescent images of fibroTUG tissues of varied fiber alignment, fiber stiffness, and post stiffness seeded with iPSC-CMs possessing a GFP-TTN reporter. Maximum contractile (b) force, (c) stress, and (d) work quantified in tissues with constant post stiffness (0.41 N/m) with varied fiber alignment and fiber stiffness (n ≥ 11 tissues). (e) Sarcomere alignment deviation and (f) sarcomere density quantified in tissues with constant post stiffness (0.41 N/m) and varying fiber alignment and fiber stiffness (n ≥ 10 tissues). Maximum contractile (g) force, (h) stress, and (i) work quantified in tissues with constant matrix stiffness (0.68 kPa) with varied fiber alignment and post stiffness (n ≥ 11). (j) Sarcomere alignment deviation and (k) sarcomere density quantified in tissues with constant matrix stiffness (0.68 kPa) with varied fiber alignment and post stiffness (n ≥ 8). All data presented as mean ± std; * p < 0.05.

Regardless of matrix conditions, all tissues contracted uniaxially, as evidenced by inward post deflections (**Supp. Videos S1-S8**), similar to established rectangular micropatterned 2D and 3D tissues^17,18,21,22,26^. Examining fibroTUGs formed on aligned fibrous matrices spanning soft (0.41 N/m) posts, we noted a decrease in tissue contractile force, contractile stress, and work as a function of increasing matrix stiffness (**Fig. 2b-d; Supp. Video S1**). Exploring the effect of matrix fiber alignment (via pre-defining aligned vs. randomly oriented fibers), we noted diminished tissue contractility on soft (0.68 kPa) matrices with randomly oriented fibers as compared to matrices composed of aligned fibers of equivalent fiber stiffness (**Fig. 2b-d; Supp. Video S1,S2**). On stiffer matrices, differences in tissue stress arising from fiber alignment was less prominent, potentially indicating that iPSC-CMs may be unable to deform stiffer matrices or efficiently assemble myofibrils independent of matrix fiber alignment. We next quantified myofibril organization within these tissues and observed a decrease in sarcomere alignment on randomly oriented fiber matrices regardless of matrix stiffness, as quantified by a higher sarcomere deviation^15^ (**Fig. 2a,e,f**). Additionally, sarcomere density decreased on random matrices compared to aligned matrices across all stiffnesses tested (**Fig. 2a,e,f**). Finally, we examined the influence of tissue boundary stiffness on resulting tissue contractility by maintaining a constant stiffness of aligned matrices at 0.68 kPa while increasing post stiffness (**Fig. 2g-i; Supp. Video S3,S4**). While contractile force and stress remained constant in tissues formed between both soft (0.41 N/m) and stiff (1.2 N/m) posts, the effective work produced by tissues contracting against stiffer boundaries was greatly reduced (**Fig. 2g-i**). However, tissues formed on randomly oriented, soft fiber matrices tethered between stiff posts surprisingly revealed no differences in contractile force and stress as compared to aligned, soft matrices suspended between stiff posts (**Fig. 2g,h**). This finding may be explained by a greater relative influence of increased uniaxial workload against the stiff posts, given that these tissues also exhibited enhanced myofibril assembly and alignment compared to tissues formed on random matrices under soft boundary conditions (**Fig 2,j,k**). This surprising result suggests that iPSC-CMs contracting against stiffer boundary constraints can form aligned myofibrils regardless of the topographical alignment of the underlying matrix, as has been shown previously in 3D tissues^21,50,51^. However, as these tissues demonstrated limited fractional shortening and work, this condition may represent a diseased or supraphysiologic mechanical environment^43^.

We next explored how microenvironmental mechanics impact iPSC-CM EHT maturation. To assess structural maturation, we immunostained tissues formed in different mechanical environments for connexin-43, the predominant cardiac gap junction protein, and the myosin regulatory light chain MLC-2v, which is known to be enriched in adult ventricular CMs^7^ (**Fig. 3a-d**). Corroborating our measurements of fibroTUG contractility and myofibril assembly, tissues formed on soft (0.68 kPa), aligned matrices and soft (0.41 N/m) posts expressed the highest levels of connexin-43 and MLC-2v compared to tissues formed on stiff aligned matrices, soft non-aligned matrices, or between stiff posts (**Fig 3a-d**). Additionally, cardiac troponin T expression was also most abundantly expressed in tissues formed on soft (0.68 kPa) aligned matrices and soft (0.41 N/m) posts (**Fig. S3**).

**Figure 3:**
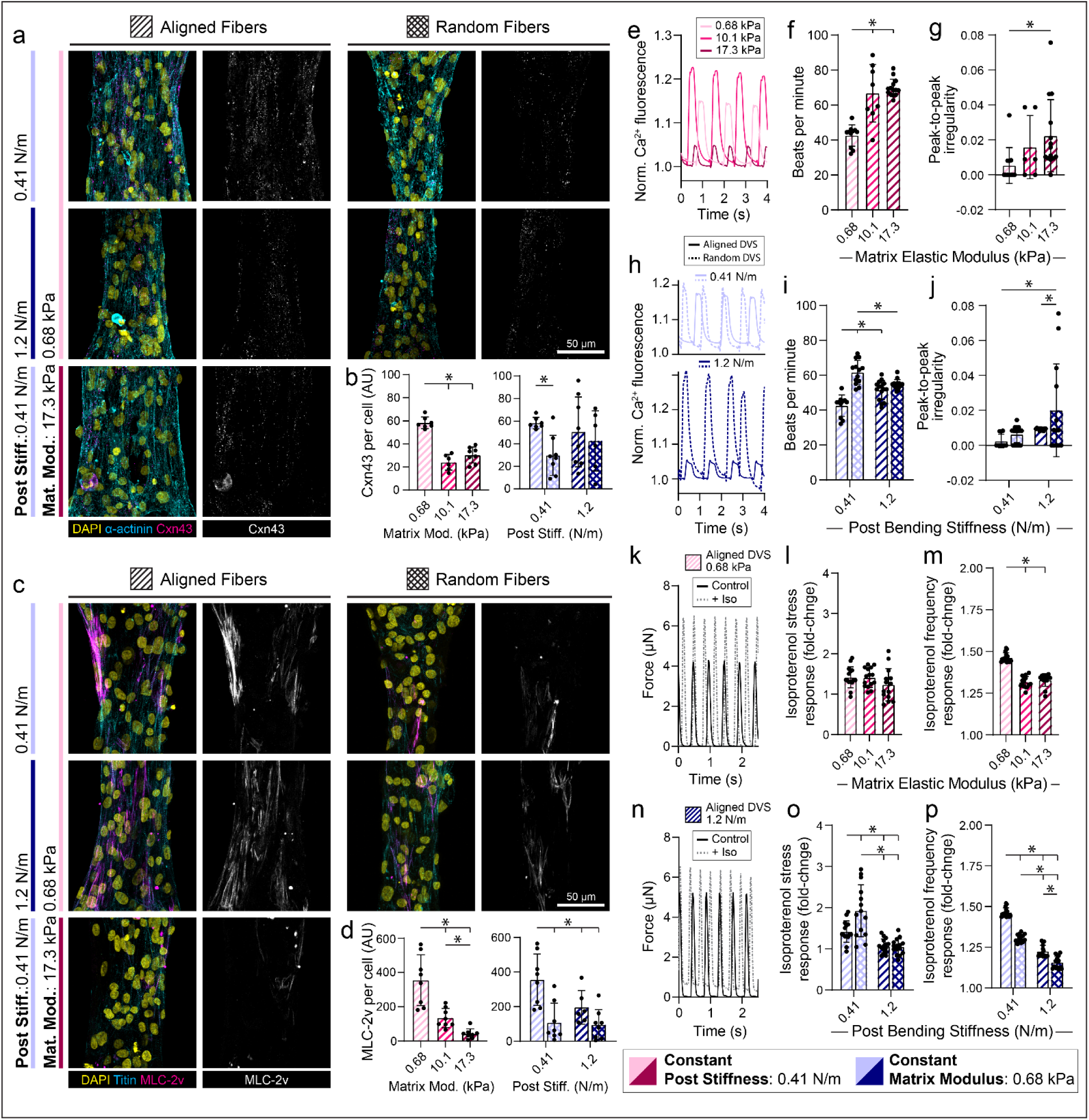
Fibrous matrix alignment, stiffness, and mechanical constraints influence iPSC-CM tissue development. (a) Confocal fluorescent images of fibroTUG tissues of varied fiber alignment, fiber stiffness, and post stiffness immunostained for α-actinin and connexin-43. (b) Quantification of connexin-43 (Cxn43) expression (n ≥ 6). (c) Confocal fluorescent images of fibroTUG tissues of varied fiber alignment, fiber stiffness, and post stiffness seeded with iPSC-CMs containing a GFPtitin reporter and immunostained for MLC-2v. (d) Quantification of MLC-2v expression (n ≥ 6). Calcium flux dynamics were analyzed, with representative flux traces shown in (e) and (h), to determine (f,i) contraction frequency (n ≥ 8) and (g,j) peak-to-peak irregularity, as quantified by the standard deviation of time interval between peaks (n ≥ 6). Contractile dynamics in response to isoproterenol treatment (10 nM) were analyzed, with representative contraction traces shown in (k) and (n), to determine the fold change in (l,o) contractile force and (m,p) contractile frequency (n ≥ 14). Blue hatching within bar plots indicates where post stiffness was held constant at 0.41 N/m in (b,d,e-g,k-m) to explore the impact of matrix alignment and post stiffness on tissue development. Pink hatching within bar plots indicates matrix stiffness was held constant at 0.68 kPa in (b,d,h-j,n-p) to explore the impact of matrix alignment and fiber stiffness on tissue development. All data presented as mean ± std; * p < 0.05.

To further corroborate these findings, we next assessed calcium handling of tissues formed with the fibroTUG platform. Briefly, tissues were incubated with a calcium sensitive dye and imaged at frame rates > 65 frames/sec. Quantification of calcium flux dynamics indicated tissues formed on soft (0.68 kPa) aligned matrices with soft (0.41 N/m) posts contracted at the lowest frequency and at the most regular intervals (in contrast to heightened peak-to-peak irregularity observed on tissues formed on stiff or random matrices, suggestive of heightened arrythmogenic activity); of note, decrease contraction frequency and regularity are both considered to be characteristic of more mature iPSC-CMs^7^ (**Fig. 3e-j; Supp. Video S5-S7**). We also treated tissues with the β-adrenergic agonist isoproterenol to further analyze tissue maturation and function (**Fig. 3k-p**). As β-adrenergic signaling plays a critical role in regulating cardiomyocyte contractility and calcium handling, robust responses to agonists of this pathway, such as increased contractile frequency and stress, are generally indicative of a more mature phenotype^7,18,21,52^. Across all conditions tested, tissues demonstrated chronotropic and inotropic responses to isoproterenol (**Fig. 3k-p; Supp. Video S8**). Importantly, isoproterenol induced a greater force-frequency response in tissues formed on soft (0.68 kPa), aligned matrices with soft (0.41 N/m) posts as compared to tissues formed on stiffer aligned matrices, soft non-aligned matrices, or stiffer posts (**Fig. 3k-p**).

These findings highlight the value of bioengineered platforms that enable orthogonal tuning of various microenvironmental mechanical inputs, as each input appeared to have unique effects on tissue structure and function. Collectively, these experiments support the claim that tissues formed on soft, aligned matrices contracting against soft boundaries resulted in the most structurally and functionally mature EHTs. Furthermore, these studies also suggest that combinations of mechanical and architectural inputs alter tissue structure and function in complex and non-intuitive ways, motivating the use of this highly controlled system to study iPSC-CM tissue maturation. For example, while randomly oriented fibers in most cases led to less contractile tissues with disorganized myofibrils, surprisingly, randomly oriented fibers spanning stiff posts yielded tissues with aligned myofibrils and heightened contractility. Despite mirroring the highest functioning tissues in terms of contractile stress and myofibril organization, however, these tissues displayed substantial contractile frequency irregularity, consistent with a pro-arrhythmic phenotype and expressed lower levels of connexin-43 and MLC-2v (**Fig. 2,3**). Leonard et al. showed that gradually increasing cantilever stiffness yields increases in contractile force to a point where the boundary stiffness reached potentially pathologic levels^43^. Additionally, increased mechanical loading of tissues has been shown to increase myofibril organization^51,53^, further supporting the fact that high boundary stiffness may be driving this phenotype.

Finally, while these studies indicate the importance of control over mechanical features and matrix architecture in driving iPSC-CM maturation, other maturation techniques such as electrical pacing and metabolic programming are likely essential in deriving tissues that more closely approach the function of healthy adult myocardium^21,54,55^. Indeed, culturing fibroTUGs formed with varying post stiffnesses in media that promotes oxidative phosphorylation yielded tissues with greater contractile force after 7 and 14 days in culture as compared to those cultured in standard media^18^ (**Fig. S2**).

### Tissue specific modeling of fibroTUG reveals altered cellular response to matrix stiffness

The ability to orthogonally tune mechanical and architectural inputs to EHT formation enables investigation into how iPSC-CMs sense and respond to distinct physical microenvironmental cues. However, a limitation of many EHT platforms (including ours) is that tissue-scale contractile force readouts are determined by measuring post deflections. While this affords a direct measure of dynamic tissue contractility, these measurements fail to capture stresses and strains at the cell and subcellular levels. This is best illustrated by an example of highly contractile CMs contracting on rigid matrices, where limited post deflections would misleadingly suggest low CM contractility. The discrepancy between tissue contractility and cell force/stress thus limits our interpretation of how iPSC-CMs may be responding to matrix alignment, matrix stiffness, or boundary constraints. To overcome this challenge, we generated tissue-specific computational models of fibroTUGs that enable quantification of the active cell stresses generated by iPSC-CMs within these tissues based on the input parameters of composite force, myofibrillar structure, and the underlying fiber structure. Detailed methods on how tissue-specific models were generated and validated can be found in Jilberto et al. (2023)^49^. Briefly, to generate these computational models, an image analysis pipeline was developed to extract key matrix and cell parameters including fiber density, alignment, and dispersion; sarcomere density and alignment; and tissue displacements and sarcomere strain from time-lapse imaging^15^ (**Fig. 4a-c**). From these metrics, a non-linear hyperelastic finite element model that accounts for tissue-specific fiber and cell mechanics was constructed. The contraction of each tissue was simulated while computing the active stress of CMs required to generate the experimentally determined contractile dynamics. By design, the model captures the contractile behavior experimentally measured at the posts (**Fig. 4d**) and the active stress curve generated by the model describes the heterogenous local contractile function that the CMs generate given the tissue-specific inputs described above (**Fig. 4e**).

**Figure 4:**
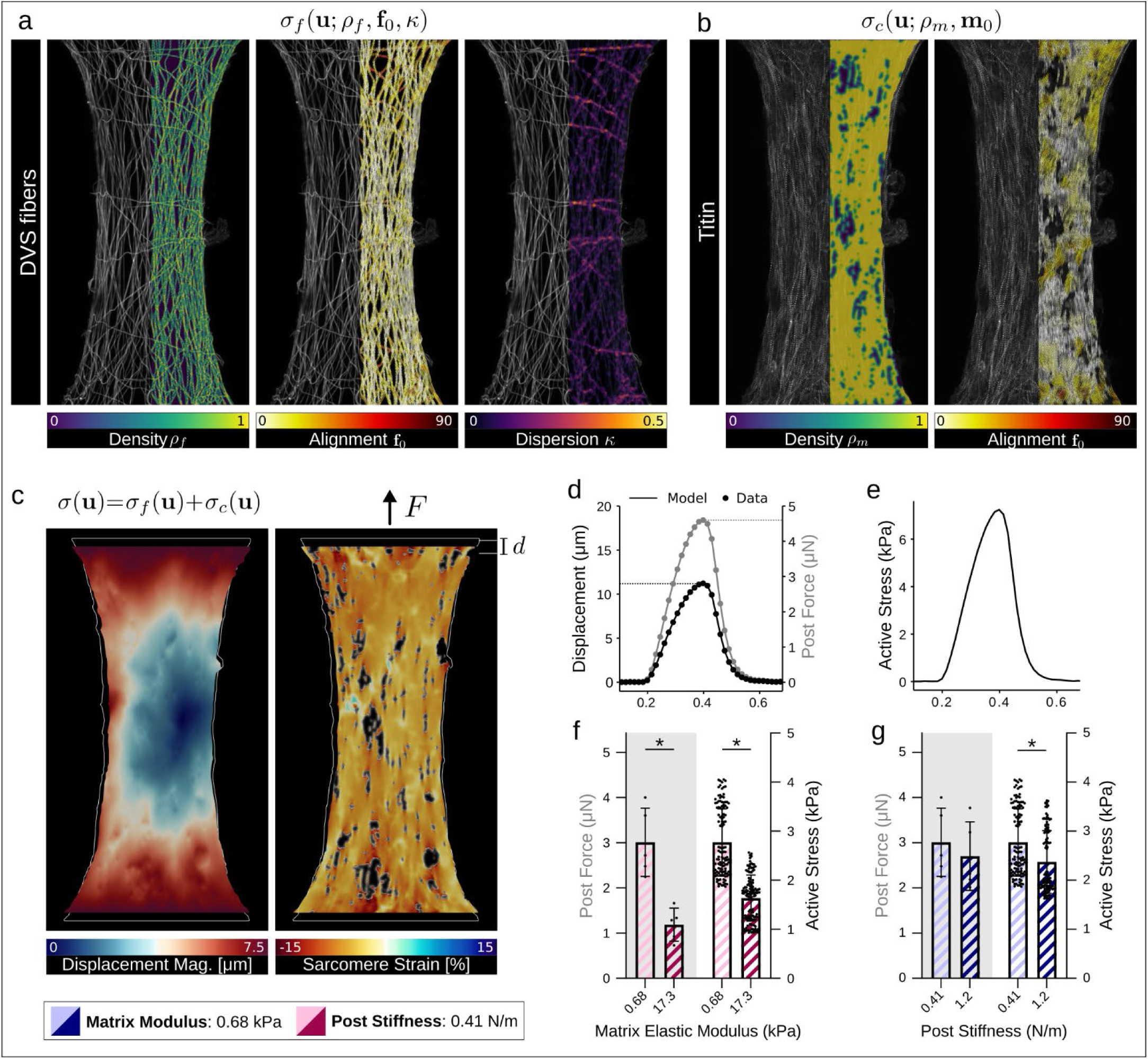
Tissue-specific computational modeling of fibroTUGs shows altered cellular contractility on matrices of varied stiffness. (a) Density, alignment, and dispersion fields characterizing the structure of the fibrous matrix. (b) Sarcomere density and alignment characterizing the structure of the myofibril network. (c) Results of the simulation for representative tissue showing inner displacement magnitude (left), and sarcomere strain (right). (d) Simulated post displacement and force time traces matching the experimental data for one simulation, where post displacement and force data are the simulation input. (e) The resulting mean active stress curve exerted by sarcomeres in the model to match the data in (d). (f-g) Post-force input (left two bars on gray background) compared with the computed active stress (right two bars on white background) for n>100 simulations with varied fiber stiffness (f) and varied post-stiffness (g). All data presented as mean ± std; * p < 0.0001 by unpaired t-tests.

In comparing tissues formed with soft vs. stiff fibers, we found that CMs on stiff matrices in fact generated lower active cell stresses than those on soft matrices (**Fig. 4f**), suggesting that a cellular mechanoresponse to matrix stiffness in part explains the observed reduced force output. The extent to which decreased post-derived force measurements on stiff matrices is a product of an adaptive cellular response as opposed to the mechanics of the fiber matrices is unknown. However, the magnitude of the decrease in post force on stiff matrices compared to soft was greater than the magnitude of the decrease in the cellular active stress on stiff matrices (**Fig. 4f**), highlighting a combination of cellular mechanosensing in addition to matrix stiffness and structure in defining tissue force measured by post deflections. Post stiffness, however, did not significantly change the relationship between tissue and active cell stresses, as expected (**Fig. 4g**). Further exploration of insights gained from the computational model are discussed in the concurrent manuscript^49^. Here, we delved deeper into the discussed result and its implications - that matrix stiffness impacts how CMs generate intercellular forces and forces applied extracellularly to the underlying matrix.

### Matrix stiffness impacts cell-ECM interactions and costamere formation

The preceding studies demonstrate that matrix stiffness significantly impacts EHT formation and function (**Fig. 2**,**3**), expression of markers associated with maturation (**Fig. 3**), and active cell stresses (**Fig. 4**). As the fibroTUG platform provides a means to probe how the mechanics of the fibrous ECM impacts tissue assembly and maturation, we next examined if cell-ECM interactions might differ in tissues formed on soft compared to stiff matrices. Previous work from Chopra et al. implicated focal adhesions (FAs) or protocostameres as critical nucleation points for sarcomere and myofibril assembly in iPSC-CMs cultured on 2D micropatterns^32^. Moreover, Fukuda et al. examined the role of vinculin, a mechanosensitive protein that localizes to cell-matrix adhesions as well as adherens junctions, in zebrafish heart development^30^. Their results indicate that mechanical strain upregulates vinculin expression, which is known to mediate myofibril maturation throughout development. We thus hypothesized that altered interactions as a function of fibrous matrix stiffness could impact FA and myofibril assembly. To test this hypothesis, we generated tissues between soft posts, on aligned soft (0.68 kPa) or stiff (17.3 kPa) matrices and assessed after 1, 3, or 7 days of culture. Immunostaining for vinculin, a marker of mechanoresponsive FAs and key regulator of cardiac development^30^, and quantification of FA size, shape, and overall abundance revealed marked stiffness-mediated differences in cell-matrix interactions during tissue assembly and maturation (**Fig. 5**). At day 1, during initial myofibril assembly, FAs were observed to colocalize with matrix fibers and the number of vinculin-rich FAs, average size of FAs, total vinculin expression, and the eccentricity of each adhesion all were significantly greater on soft matrices as compared to stiff (**Fig. 5a-e**). Further, initially formed immature myofibrils were more disorganized on stiff matrices than on soft (**Fig. 3a, S4**). At days 3 and 7, the average FA size and total FA signal, as determined by vinculin immunostaining, decreased slightly independently of matrix stiffness (**Fig. 5c,d**). At these later timepoints, vinculin localized to z-discs most prevalently in tissues formed on soft matrices as quantified by colocalization with the z-disc protein titin. This co-localization that suggests the formation of costameres to our knowledge has not been previously reported for iPSC-CM EHTs (**Fig. 5a,e-g**). In native myocardium, costameres physically link myofibrils to the surrounding ECM at each z-disc, enabling force transmission to adjacent tissue^29,41,42^. These structures are known to play a critical role in regulating myocardial contractile function and their formation during development may be regulated in part by mechanical strain within the tissue^30,31^. We also observed an increase over time in the expression of the β1D integrin splice isoform uniquely in soft matrices (**Fig. S5**). Integrin β1D is specific to cardiac and striated muscle cells and has previously been associated with cardiac maturation ^56–58^. Taken together, the formation of costameres and increased expression of β1D integrin may indicate a more mature CM adhesive state regulated by matrix mechanics.

**Figure 5:**
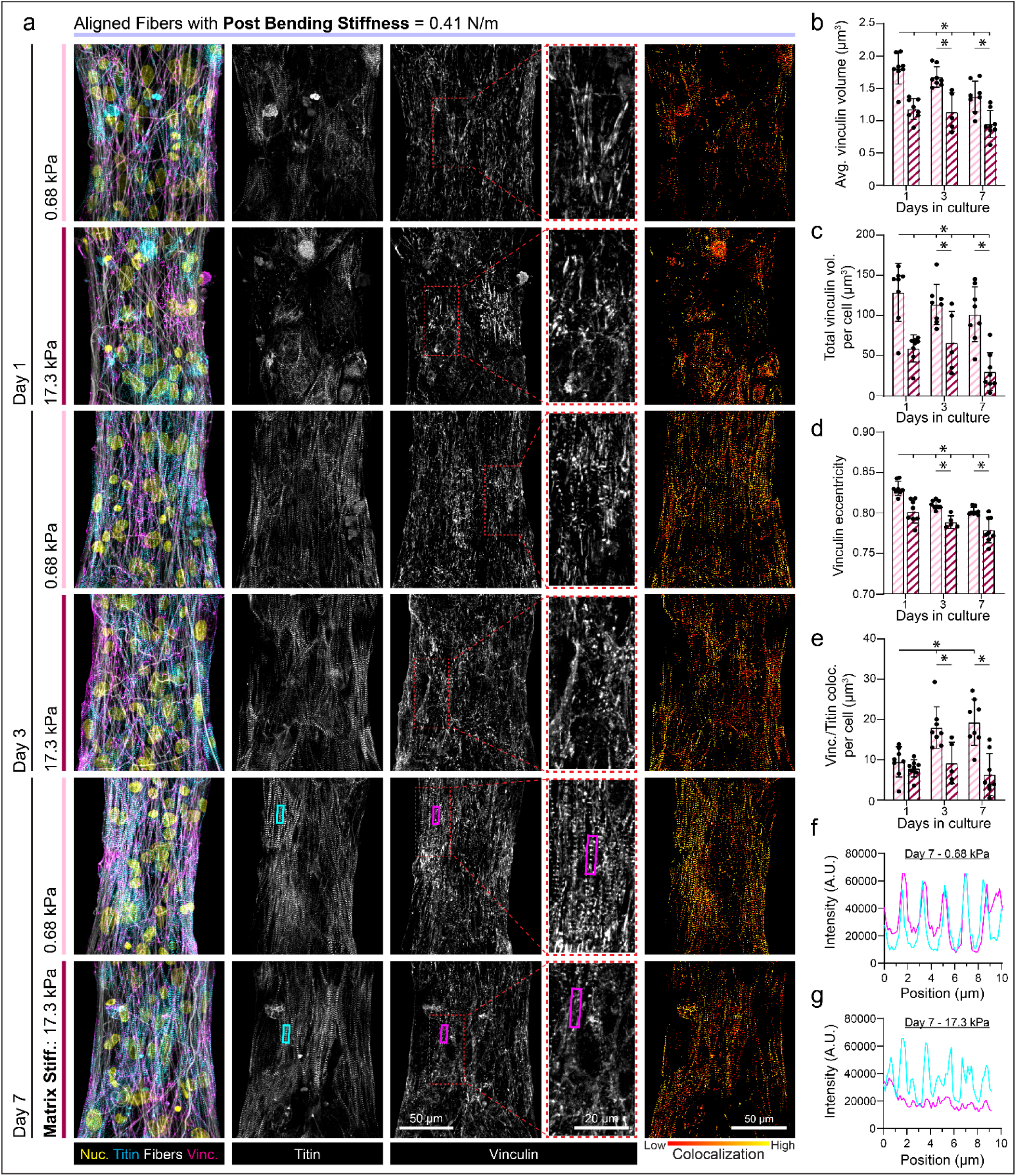
Matrix mechanics influence costamere formation which regulates myofibril assembly and maturation. (a) Confocal fluorescent images of fibroTUG tissues fixed at day 1, 3 and 7 post seeding on either soft (0.68 kPa) or stiff (17.1 kPa) aligned fiber matrices (post stiffness was held constant at 0.41 N/m). (b) Average vinculin volume, (c) total vinculin volume, (d) and vinculin eccentricity were quantified from the fluorescent images of immunostained vinculin (n ≥ 5). (e) Costamere formation was assessed by quantifying vinculin colocalization with titin. Colocalization of vinculin and titin on day 7 was visualized via fluorescence intensity plots of titin (cyan) and vinculin (magenta) on (f) 0.68 kPa matrices and (g) 17.1 kPa matrices from lines drawn along the major axis of regions indicated by the rectangles overlayed on images in panel a. All data presented as mean ± std; * p < 0.05.

As vinculin also localizes to adherens junctions, we co-immunostained tissues fixed at days 1, 3, and 7 for N-cadherin and vinculin to examine whether N-cadherin (N-cad) and vinculin co-localization to the intercalated disc was also influenced by matrix stiffness (**Fig. 6a-d**). Total N-cadherin expression increased comparably over culture time in tissues formed on both soft and stiff matrices (**Fig. 6a,b**). Expression of desmoplakin, a key desmosomal protein, also increased similarly with time on soft and stiff matrices (**Fig. S6**). Localization of vinculin to N-cadherin, however, was greatest on soft matrices at days 3 and 7, potentially indicating the formation of more robust and mechanically engaged intercalated discs (**Fig. 6 a,c**). When normalizing the intensity of vinculin at intercalated discs to costameric vinculin, differences between the two matrix conditions were not apparent, indicating that vinculin expression is upregulated in both locations in the more contractile tissues formed on soft matrices (**Fig. 6d**). This idea is supported by findings from Fukuda et al. that indicate vinculin localization to both costameres and cell-cell junctions is upregulated in CMs that experience mechanical strain during development^30^.

**Figure 6:**
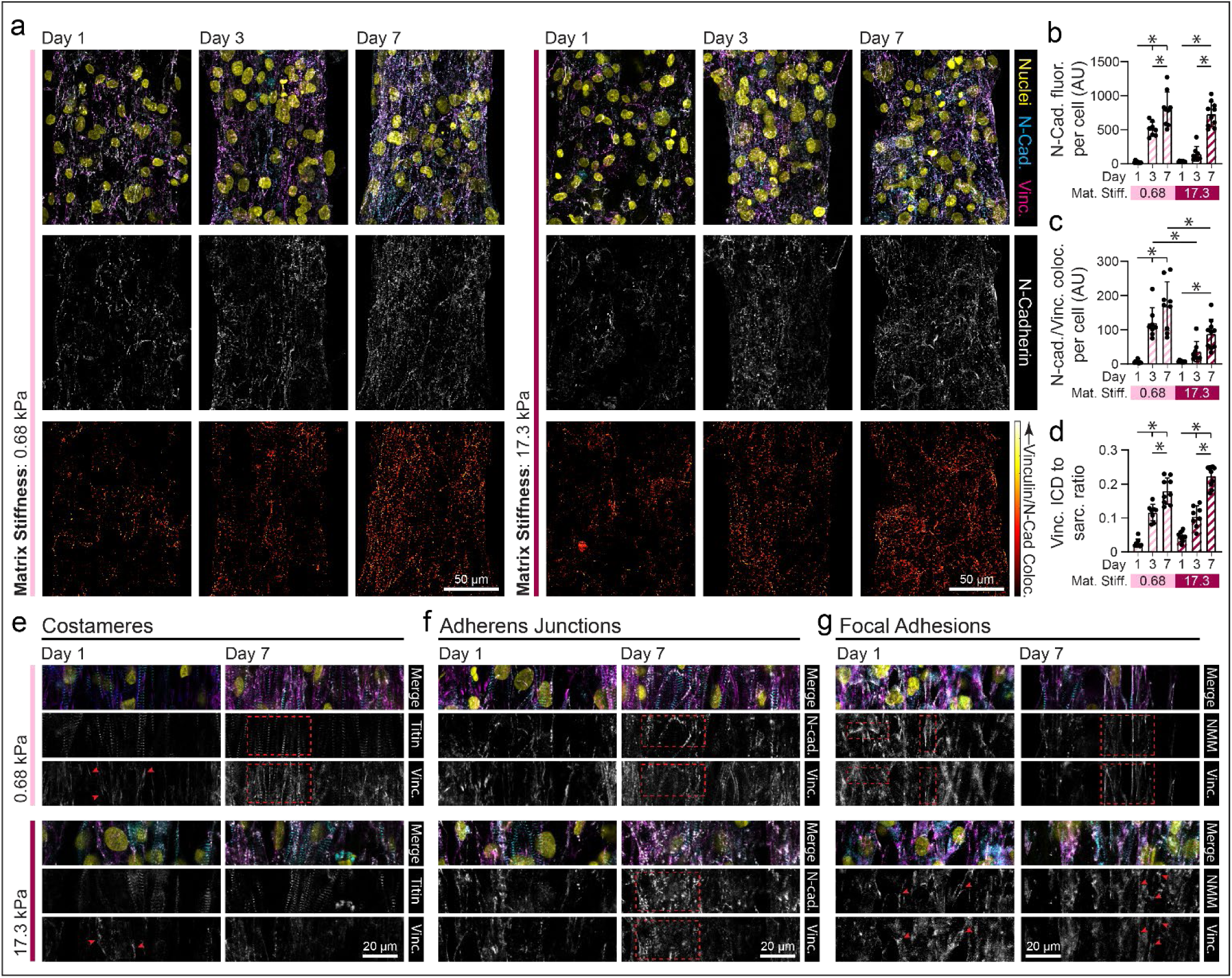
Microenvironmental mechanics regulate vinculin localization to costameres, adherens junctions and focal adhesions. (a) Confocal fluorescent images of fibroTUG tissues fixed at day 1, 3 and 7 after seeding on either soft (0.68 kPa) or stiff (17.1 kPa) aligned fiber matrices (post stiffness was held constant at 0.41 N/m). (b) N-cadherin (N-Cad) fluorescence was quantified from the fluorescent images of immunostained N-cadherin (n ≥ 8). (c) Vinculin localization to adherens junctions was quantified by analyzing vinculin and N-cadherin immunostained images. (d) Ratio of vinculin that localized to intercalated discs (ICD) to vinculin localized to costameres or sarcomere z-discs. Representative confocal fluorescent images of vinculin localization at days 1 or 7 to (e) costameres (i.e., vinculin colocalization to titin), (f) adherens junctions (i.e., vinculin colocalization to N-cadherin), and (g) focal adhesions (i.e., vinculin colocalization to NMM-IIB) not associated with either N-cadherin or titin staining, as indicated by red arrow heads and boxes. All data presented as mean ± std; * p < 0.05.

Using high resolution imaging, we identified three distinct locations to which vinculin localizes in fibroTUG tissues: 1) FAs, most prevalent upon initial cell adhesion and during myofibril assembly (day 1), 2) intercalated discs, and 3) costameres, which were evident by day 7 following myofibril formation^41^ (**Fig. 6e-g**). Of note, vinculin localized to z-discs to form costameres preferentially in soft matrices by day 7, suggesting myofibril maturation may be associated with the formation of these critical cell-ECM adhesions (**Fig. 6e**). Vinculin was present at N-cadherin-rich adherens junctions in both soft and stiff matrices by day 7 (**Fig. 6f**), despite the lower amount of N-cadherin colocalization with vinculin observed in stiff tissues more broadly (**Fig. 6c**). Finally, we observed vinculin localization to FAs distinct from z-discs based on the lack of titin and instead co-localization with non-muscle myosin-IIB (NMM-IIB) (**Fig. 6g**). FAs were particularly prominent in assembling tissues on day 1 and more commonly observed on stiff matrices across all time points (**Fig. 6g**), in line with decreased vinculin localization to titin noted on stiff matrices (**Fig. 5e**). Furthermore, these complexes were larger and more elongated on soft matrices than stiff matrices at day 1, suggesting more robust adhesion to soft matrices, as previously described (**Fig. 6g, 5d**). These three vinculin-enriched adhesive structures were first observed by Simpson and colleagues in adult feline CM cultures^41^. Their results suggest that contractile behavior and the formation of cell-cell junctions regulate vinculin distribution and potentially myofibril assembly^32,38,41,42^. Our observations indicate that cellular mechanosensing of the surrounding ECM drives the expression and localization of vinculin to distinct cellular domains within iPSC-CMs and further supports the role that contractile activity plays in regulating vinculin distribution (**Fig 5**, **6**).

### Tissue contractility drives myofibril maturation and costamere formation in soft matrices

In the preceding studies, soft (0.68 kPa), aligned fibrous matrices yielded the most contractile tissues and localization of vinculin to costameres, an adhesive phenotype not previously reported in EHTs derived from iPSC-CMs (**Fig. 2**,**5**,**6**). As prior studies implicated myosin contractility as being critical for myofibril assembly and maturation^18,32,37,59^, we next examined if myosin contractility also plays a role in regulating myofibril stability and costamere formation. iPSC-CMs seeded on soft matrices were treated with blebbistatin (50 µM) beginning at day 3, an inhibitor of both non-muscle myosins and cardiac myosins, or mavacamten (500 nM), a cardiac myosin specific inhibitor, to test whether myosin contractility is required for myofibril maturation and concomitant vinculin localization to costameres (**Fig. 7**). Comparisons were made to tissues formed on stiff matrix conditions, which exhibited less costameric vinculin and overall lower myofibril density (**Fig. 5**). Tissues were analyzed on day 8 to assess diastolic stress, vinculin localization, and myofibril assembly. In only untreated tissues on soft matrices, we observed a decrease in diastolic tissue length implying enhanced tissue contractility (**Fig. 7b**). In contrast, blebbistatin and mavacamten treated tissues both increased in length, indicating relaxation and lower diastolic stress (**Fig. 7c-d**). Further, the number of vinculin-enriched adhesions, total vinculin expression, and eccentricity of adhesive structures decreased in tissues treated with both blebbistatin and mavacamten from days 3-8, implying disrupted maturation of cell-cell and cell-ECM adhesions (**Fig. 7e-g**). FA organization and vinculin localization in treated tissues formed on soft matrices were similar to that of untreated stiff matrix tissues, suggesting that diminished CM contractility on stiff matrices may limit FA maturation (**Fig. 7e-g**). No effect was observed when treating tissues from only days 7-8, most likely due to the shorter treatment duration that was not long enough to allow significant disassembly of cell-ECM adhesions and myofibrils which may happen more gradually (**Fig. S7**). Additionally, myofibril organization and density, along with non-muscle myosin IIB expression, decreased upon myosin inhibition, supporting a role for actomyosin contractility in myofibril maintenance (**Fig. 7h,i, S8**). Finally, vinculin localization to z-discs decreased in treated tissues, implying that actomyosin contractility is critical for the formation of costameres (**Fig. 7j**).

**Figure 7:**
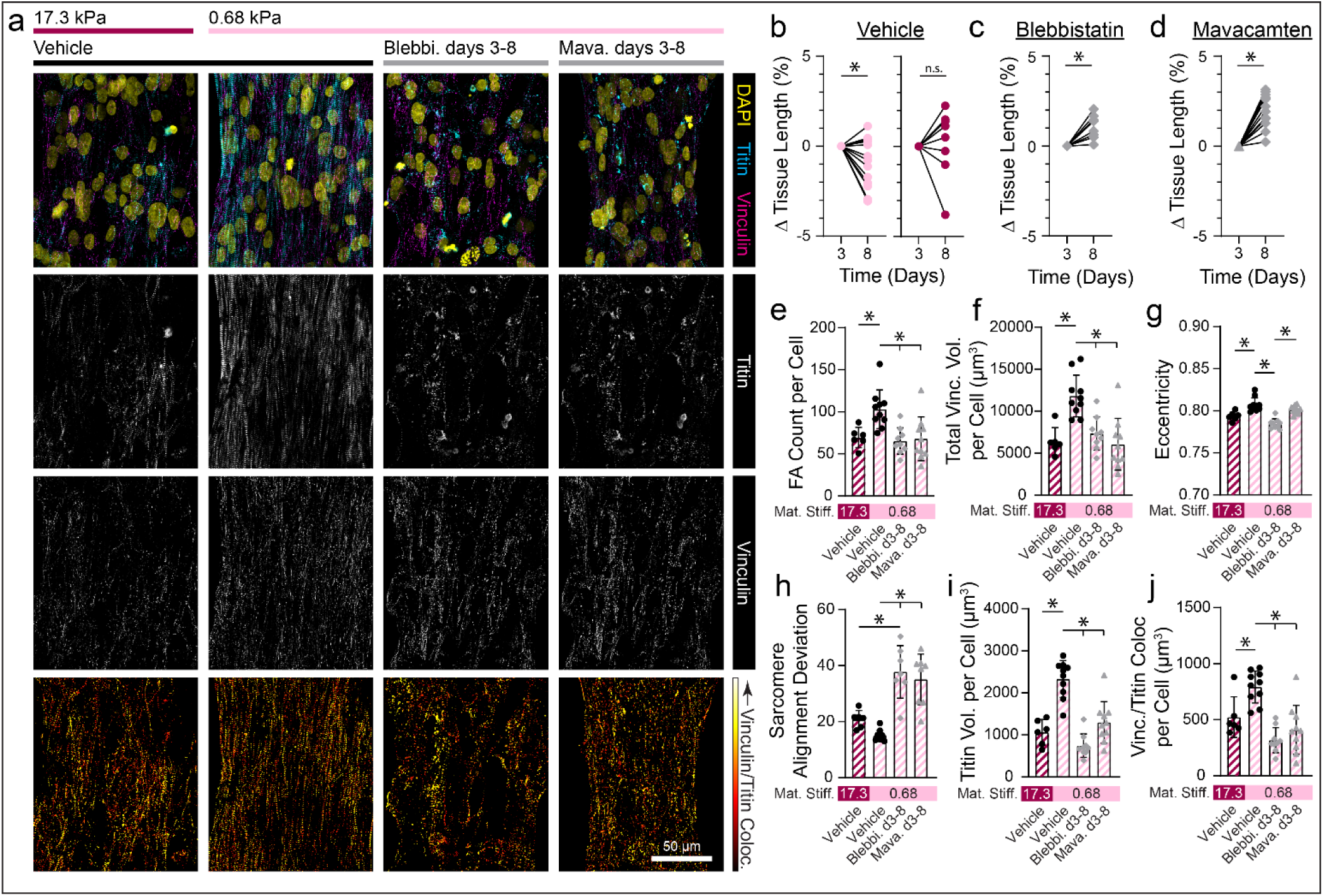
Tissue contractility drives the maturation and maintenance of myofibrils and costameres. (a) Confocal fluorescent images of fibroTUG tissues treated with blebbistatin (50µM) or mavacamten (500 nM). (b) Diastolic tissue length on day 3 and 8 of tissues seeded on soft (0.68 kPa) and stiff (17.1 kPa) aligned fiber matrices (post stiffness was held constant at 0.41 N/m) without treatment with the contractile inhibitors. (c,d) Diastolic tissue length of tissues seeded on soft matrices on day 3 before treatment with contractile inhibitors, (c) blebbistatin or (d) mavacamten, and day 8 after 5 days of treatment (n ≥ 8). (e) Focal adhesion count, (f) vinculin volume per cell, and (g) focal adhesion eccentricity were quantified from the fluorescent images of immunostained vinculin (n ≥ 6). (h) Sarcomere alignment deviation and (i) titin volume per cell quantified from fluorescent images of titin-GFP reporter (n ≥ 6). Vinculin colocalization with titin per cell quantified from titin and vinculin images (n ≥ 6). All data presented as mean ± std; * p < 0.05.

Variants in β-cardiac myosin (MYH7) and other regulators of contractility are associated with hypertrophic cardiomyopathies (HCM) and dilated cardiomyopathies (DCM). In fact, nearly a third of all known HCM variants arise from mutations in MYH7, in line with our findings that myosin driven CM contractility is critical for myofibril assembly, maturation, and overall structural integrity^34,35^. Additionally, many costameric proteins including vinculin and filamin C, are strongly implicated in dilated cardiomyopathies^34,60,61^. Studying the impact of these mutations on tissue function requires accurate models of the myocardium that possess adult-like function and structure^7^. The fibroTUG platform provides the mechanical control and ECM-like architecture of fibrous matrices necessary to driving a mature cell-adhesive phenotype that may be critical to a deeper understanding of the mechanisms of such diseases.

### Costamere formation is associated with more mature myofibrils

As costameres are the sole mediator of CM-ECM adhesion in the mature adult myocardium, we next tested the hypothesis that MLC-2v expression directly correlates with costamere formation (**Fig. 8**). Shared expression of costameric vinculin and MLC-2v could suggest that robust cell adhesion to the matrix along the myofibril facilitates myofibril maturation via recruitment of the more mature myosin light chain isoform. We seeded iPSC-CMs containing a GFP-titin reporter on both soft and stiff aligned matrices tethered between soft posts and immunostained for MLC-2v and vinculin after 7 days of culture (**Fig. 8**). As before, we observe increased MLC-2v expression and vinculin localization to z-discs on soft matrices. In contrast, iPSC-CMs on stiff matrices revealed significantly lower MLC-2v expression (**Fig. 8a,b**). Interestingly, examining the ratio of MLC-2v colocalized with vinculin-rich costameres, we found that the percentage is high in both soft and stiff matrices, despite decreased overall expression of MLC-2v in stiff matrices, indicating a relationship between myofibril maturation and costamere formation (**Fig. 8c**). To confirm this quantification, we segmented the tissue into subsections comparable to the area inhabited by a single spread CM. For each of these subregions, MLC-2v expression was plotted against costameric vinculin and fit to a linear regression model (**Fig. 8d,e**). On both soft and stiff matrices, correlation between these two metrics was positive and significant; however, a higher linear regression slope was noted for tissues formed on soft matrices, indicating that the highest level of both costameric vinculin MLC-2v was attained in tissues on soft matrices (**Fig. 8d,e**). Taken together, these data indicate that the formation of costameres is coupled with myofilament maturation, though costamere formation does not necessarily lead to myofibrillar maturation. This highlights the critical relationship between CM cell-matrix adhesion and myofibril maturation that hand in hand define CM and overall tissue contractility.

**Figure 8:**
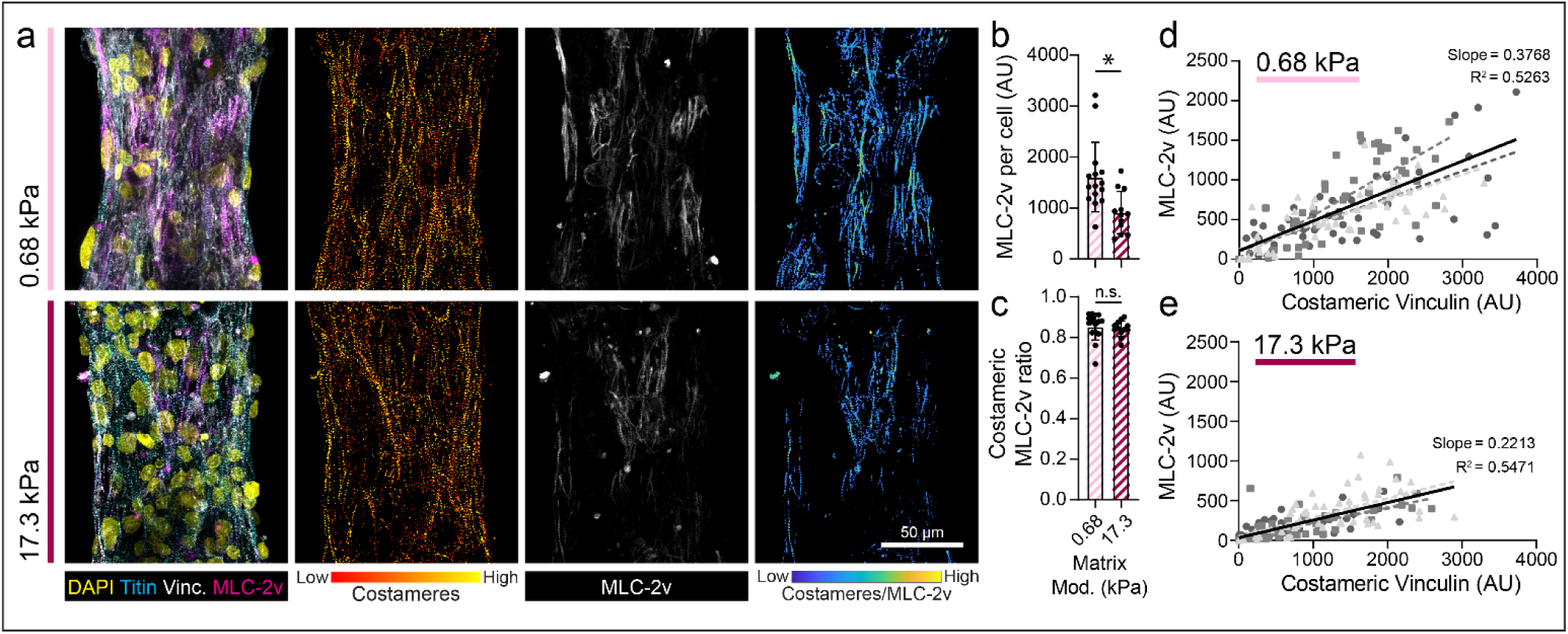
Costamere formation is associated with more mature myofibrils. (a) Confocal fluorescent images of fibroTUG tissues fixed at day 7 after seeding on either soft (0.68 kPa) or stiff (17.1 kPa) aligned fiber matrices (post stiffness was held constant at 0.41 N/m). (b) Quantification of MLC-2v expression per cell (n ≥ 12). (c) Ratio of MLC-2v expressing myofibrils that are associated with robust costamere formation (n ≥ 12). For tissues formed on (d) soft and (e) stiff matrices, correlation between MLC-2v expression and costameric vinculin was assessed by segmenting tissues into regions equal to roughly the size of one CM and MLC-2v and costameric vinculin expression was quantified within each of these regions. Each point on the plot represents a single subregion on one of three representative tissues. Linear regressions for data from each individual tissue are indicated by different colored dashed lines while linear regression of the data from these three tissues pooled together is indicated by the solid black line. The slope and R^2^ value for this pooled data are also noted in the plots. All data presented as mean ± std; * p < 0.05.

## DISCUSSION

Studying the impact of biophysical cues on iPSC-CM tissue assembly and maturation despite many recent advances remains challenging. Current 2D models provide an excellent testbed for certain applications, but typically cannot recapitulate native tissue mechanics and architecture. In contrast, 3D constructs more closely mimic native environmental conditions and can more effectively drive tissue maturation; however, isolating the impact of tissue-relevant mechanical and architectural cues on CMs in these models is intractable^8,9^. Here, we created a 2.5D tunable iPSC-CM tissue platform composed of mechanically tunable synthetic fiber matrices suspended between two elastomeric posts, enabling the study of key physical microenvironmental inputs to tissue assembly and function: tissue boundary constraints, matrix fiber alignment, and matrix stiffness.

Unlike other approaches to engineering cardiac tissues, the orthogonal tunability of the fibroTUG cardiac microtissue platform allowed for careful dissection of how individual biophysical cues impact CM organization and function. Furthermore, the formation of thin tissues (∼15 µm) between elastomeric posts permits comprehensive analysis of both contractile function and structural development of the tissues. Our studies exploring the influence of fiber stiffness could provide insight into how changes in ECM mechanics during cardiac development or disease progression may impact cardiac tissue organization and function. Alternatively, matrix alignment is known to play a critical role in tissue function where disorganized fibrotic ECM deposited due to myocardial injury has been suggested to contribute to abnormal myocardial biomechanics and disease-associated signaling^10,62^. Finally, control over post stiffness to define tissue boundary constraints may reflect tissue remodeling associated with altered cardiac afterload, which is known to alter heart function in various contexts^35,43,51,53,63^.

Importantly, many cardiac microtissue platforms require the inclusion of stromal cells for tissue assembly and compaction^8,21–24,64,65^. However, their random distribution amongst contractile CMs does not accurately recapitulate the organization of the native myocardium where CMs and stromal cells are segregated^10,11,13^, and furthermore confounds clear insights into how various biophysical cues impact specifically CM development and maturation. Additionally, these tissues typically exhibit limited stability over time in culture due to stromal cell proliferation, contraction, and tissue delamination^7,8^. In our fibroTUG platform, predefining matrix properties before seeding enables the assembly of iPSC-CM tissues without the addition of stromal cells.

We found that tissues formed on soft (0.68 kPa), aligned matrices suspended between soft (0.41 N/m) posts were the most mature, as evidenced by myofibril density and contractility (**Fig 2**,**3**). Tissue maturity was assessed via quantification of structural development and calcium handling. We found that each mechanical input has distinct effects on tissue assembly, maturation, and contractility, suggesting that different mechanosensing mechanisms may be involved in responding to various mechanical signals. For example, altered contractile function of tissues formed in aligned vs. random fiber matrices is likely a result of disrupted myofibril organization. Contractility of CMs on soft vs. stiff fiber matrices is in part defined by differential mechanosensing and CM-ECM interactions that lead to changes in cellular active stress^49^. Further, while increasing post stiffness did not impact cellular active stress or post-force measurements, stiff boundary constraints appear to be sufficient in driving myofibril alignment regardless of matrix organization, despite low levels of fractional shortening.

The integration of finite element models of our fibroTUG platform provided further insights into how iPSC-CMs respond to the specific mechanical perturbations (**Fig. 4**). Tissue-specific modeling enabled precise analysis of how matrix architecture and mechanics impact iPSC-CM contractile function^49^. Specifically, we found that differences in tissue contractility as measured by elastomeric posts were not only caused by changes in matrix deformability, but also a cellular contractile response to changes in matrix stiffness (**Fig. 4**). Further supporting this claim, cell adhesions, and more specifically the formation of costameres, proved sensitive to matrix stiffness, implying that cellular mechanosensing dictates the formation of these structures and thus defines how tissues assembly and mature (**Fig. 5**).

Fiber alignment plays a key role in the development of fibroTUG tissues, and previously developed techniques including electrospun scaffolds, nano- and micro-grooved surfaces, or micropatterning have been shown to drive alignment and subsequent maturation of iPSC-CMs ^15–19^. Additionally, many groups have explored how matrix stiffness impacts CM maturation and contractile function, culminating in the observation that CMs contract most robustly on hydrogels of physiologic stiffness (∼8-10 kPa)^17,46^. In our studies, soft, fibrous matrices (0.68 kPa) that more closely mimic the stiffness of fetal heart tissue yielded the most contractile tissues^46^. This discrepancy could be explained by the distinct architecture and local mechanics of fibrous matrices as compared to isotropic, continuum-like hydrogel or elastomer surfaces that lack discrete fibrous structure and topography^15^. Furthermore, in our platform, tissues formed on soft, aligned matrices with soft boundary conditions displayed the highest levels of fractional shortening, a key regulator of iPSC-CM maturation ^18,51,66^. Caution should be taken, however, when attempting to compare elastic moduli of hydrogels and other *in vitro* culture platforms with the modulus of native tissues as characterization techniques vary widely across settings. Additionally, as highlighted herein, cells sense and respond to many mechanical properties of a material that are not captured in a simple elastic modulus measurement.

Our results may inform the design of larger-scale tissues patches and translatable regenerative therapies, where interactions between iPSC-CMs and biomaterials are likely critical to the proper assembly of functional, mature myocardial syncytia. Future work translating the identified optimal mechanical parameters to fully 3D tissue constructs with therapeutic potential will be essential. Furthermore, as disruptions in myocardial mechanics are a hallmark of many forms of cardiac disease, the high mechanical tunability of the fibroTUG platform could enable key insights into the mechanisms of disease and facilitate the development of better treatment options.

## ACKNOWLEDGEMENTS

S.J.D., J.J., M.E.J. C.S.C., H.K., E.L., A.S.H., D.A.N., and B.M.B acknowledge financial support from the CELL-MET Engineering Research Center (NSF EEC-1647837). S.J.D., A.S.H., and B.M.B. acknowledge financial support from the NSF (2033654). C.D.D. acknowledges financial support from the NSF Graduate Research Fellowship Program (DGE1256260). S.J.D acknowledges financial support from the NIH T32-DE007057 and T32-HL125242.

## AUTHOR CONTRIBUTIONS

Conceptualization—S.J.D. and B.M.B.; methodology—S.J.D., J.J., A.E.S., D.D.H., C.D.D., H.B., E.L., A.S.H., D.A.N., and B.M.B.; investigation— S.J.D., J.J., A.E.S., D.D.H., J.L., C.D.D., A.C., M.E.J. and H.K.; writing— S.J.D. and B.M.B.; writing and review/editing—S.J.D., J.J., D.D.H., M.E.J., H.B., C.S.C., E.L., A.S.H., D.A.N., and B.M.B..; funding— S.J.D., C.S.C., E.L., A.S.H., D.A.N. and B.M.B.; supervision—C.S.C., E.L., A.S.H., D.A.N., and B.M.B.

## METHODS

### Reagents

All reagents were purchased from Sigma Aldrich and used as received, unless otherwise stated.

### Elastomeric cantilever array fabrication

Arrays of poly(dimethylsiloxane) (PDMS; Dow Silicones Corporation, Midland, MI) postswere fabricated by soft lithography as previously described [cite DexVS paper and bdon nat mat]. Briefly, silicon wafer masters possessing SU-8 photoresist (Microchem, Westborough, MA) were produced by standard photolithography and used to generate PDMS stamps. Following silanization with trichloro(1H,1H,2H,2H-perfluorooctyl)silane, stamps were used to emboss uncured PDMS onto oxygen plasma-treated coverslips. Cantilever arrays were methacrylated with vapor-phase silanization of 3-(trimethoxysilyl)propyl methacrylate in a vacuum oven at 60 °C for at least 6 h to promote fiber adhesion to PDMS.

### DVS fiber matrix fabrication

DVS polymer was synthesized as previously described by our lab^44^. Briefly, dextran was reacted with divinyl sulfone and the product was dialyzed and lyophilized. For electrospinning, DVS was dissolved at 0.7 g mL^-1^ in a 1:1 mixture of milli-Q water and dimethylformamide with 0.6% (w/v) lithium phenyl-2,4,6-trimethylbenzoylphosphinate (LAP; Colorado Photopolymer Solutions) photoinitiator, 2.5% (v/v) methacrylated rhodamine (25 mM; Polysciences, Inc., Warrington, PA), and 5.0% (v/v) glycidyl methacrylate. This solution was electrospun on coverslips containing microfabricated cantilever arrays affixed to a custom-built rotating mandrel with a hexagonal geometry driven by an AC motor with controllable speed^15^. Electrospinning was conducted in an environmental chamber at 35% humidity with a flow rate of 0.2 ml h^-1^, voltage of 7.0 kV, and a gap distance of ∼5 cm to the grounded mandrel. After collection, fibers were stabilized by primary crosslinking under UV (100 mW cm^-2^) through a microfabricated photomask for 20 s, such that only the fibers suspended in the region spanning two posts would be crosslinked. Upon hydration, uncrosslinked fibers were dissolved away leaving isolated suspended microtissues adhered to the posts. Fiber matrices were subsequently placed in LAP photoinitiator solutions of varying concentrations and exposed again to UV (100 mW cm^-2^) for 20 s to tune fiber stiffness and sterilize substrates.

Matrices were functionalized with cell adhesive peptides cyclized [Arg-Gly-Asp-D-Phe-Lys(Cys)] (cRGD; Peptides International) via Michael-Type addition to available vinyl sulfone groups. Peptides were dissolved at 200 μM in milli-Q water containing HEPES (50 mM), phenol red (10 μg mL^-1^), and 1 M NaOH to bring the pH to 8.0. A volume of 150 µL was added to each substrate and incubated at room temperature for 30 min.

### Mechanical characterization

PDMS cantilever mechanics were characterized by deflecting individual posts with a ∼100µm diameter tungsten rod of known elastic modulus (**Fig. S1a-c**). Bending stiffness was calculated by measuring the cantilever deflection and the force applied by the tungsten rod using custom Matlab scripts. Bending stiffness was approximated using the following equations:

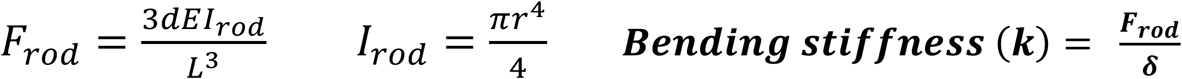

where d = rod deflection, E = elastic modulus of rod, L = length of the rod, r = radius of the rod, δ = cantilever deflection, F_rod_ = force applied by the rod, and I_rod_ = moment of inertia of the rod.

Matrix modulus was determined by pressing a microfabricated SU8 20 x 250 μm rectangle across the center of the fiber matrices to apply a tension on the matrix (**Fig. S1d-f**). Using custom Matlab scripts, matrix modulus was extrapolated from measured cantilever deflection. Matrix modulus was approximated using the following equation:

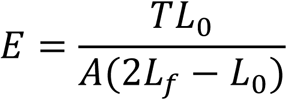

where E = elastic modulus of fiber matrix, T = tension in the matrix, A = area of the fiber matrix, L_0_ = initial length of the fibers, and L_f_ = half the final length of the fiber matrix. (More details on this calculation can be found in **Supplemental Figure 1f**.)

### iPSC culture and iPSC-CM differentiation

Induced pluripotent stem cells containing a GFP-titin reporter^67^ (PGP1; gift from the Seidman Lab) or GFP-DSP reporter (WTC; Allen Institute AICS-0017 cl.6) were cultured in mTeSR1 media (StemCell Technologies) on Matrigel (Corning) coated tissue culture plastic and differentiated via temporal Wnt modulation as previously described^4,5^. Briefly, differentiation was initiated when iPSCs reached 90% confluency in RPMI 1640 media supplemented with B27 minus insulin on day 0 with the addition of 12 µM CHIR99021 for 24 hours. On day 3, CDM3 media containing 5 uM IWP4 on day 3 for 48 hours. Retinol inhibitor BMS 453 (Cayman Chemical, 1 µM) was also added for days 3-6 to minimize atrial lineage differentiation^18,68^. Cultures were then maintained in CDM3 media until contractions began between day 8 and 10. iPSC-CMs cultures were then transferred to RPMI 1640 media lacking glucose and glutamine (Captivate Bio) supplemented with 4 mM DL-lactate, 500 ug/mL human serum albumin (Sciencell Research Labs), and 213 ug/mL L-ascorbic acid 2-phosphate trisodium salt on day 11 for 4 days. Following purification, iPSC-CMs were replated as monolayers (300,000 cells/cm^2^) on growth factor reduced Matrigel (Corning) in RPMI 1640 media supplemented with B27 for 7 additional days before seeding into tissues.

### Microtissue seeding and culture

Due to the suspended nature of the fibrous matrices, it was determined that iPSC-CMs must directly land on the top of the matrices by patterning the cells through a microfabricated cell seeding mask to prevent cells from “rolling” off the tops of the matrices and landing on the substrate beneath. Additionally, this technique significantly reduced the number of iPSC-CMs needed to efficiently seed all the tissue on the substrate. To fabricate the seeding mask, 3D printed molds were designed in SolidWorks and printed via stereolithography (Protolabs). PDMS (1:10 crosslinker:base ratio) devices were replica cast from these molds. Prior to seeding the tissues, cell seeding masks were plasma treated for 5 minutes to generate highly hydrophilic surfaces that allow water to wick through the small microtissue scale wells in the mask. Vacuum grease was then applied to the edge of the seeding mask to ensure a watertight seal between the seeding mask and the substate prior to placing the mask on the tissue array substrate such that the holes in the mask sit directly above each suspended fiber matrix.

After aligning the seeding masks, iPSC-CMs were dissociated by 0.25% Trypsin-EDTA (Gibco) with 5% (v/v) Liberase for 5 min, stopped by an equal volume of 20% FBS/1 mM EDTA/PBS. Cells were triturated by gently pipetting with a p1000 pipette eight times to obtain a near single-cell suspension and centrifuged (200 g, 4 min). iPSC-CMs were resuspended in replating media (RPMI plus B27 supplement with 2% FBS and 5µM Y-27632 (Santa Cruz Biotechnology)) and 125,000 cells were seeded per fibroTUG substrate through the top of the cell seeding mask in approximately 200 µL of media. Cultures were then moved to the incubator and left undisturbed overnight to allow the iPSC-CMs to attach to the tissues before removing the seeding masks. Cultures were maintained in RPMI media plus B27 supplement and replenished every other day for the duration of studies. All studies were carried out for 7 days unless otherwise specified.

### Contractile force analysis

Time-lapse videos of the microtissue’s spontaneous contractions were acquired at 65 Hz on Zeiss LSM800 equipped with an Axiocam 503 camera while maintaining a temperature of 37 ℃ and 5% CO2. Maximum contractile force, contractile stress, contraction kinetics, and contraction frequency were calculated using a custom Matlab script based on the deflection of the posts and the measured post bending stiffness, as described previously^22^. For isoproterenol challenge, the same tissues were imaged prior to the addition of 10nM isoproterenol and again 30 minutes after the addition of the drug for comparison.

### Calcium imaging

Calcium handling analysis was performed by incubating cells for 1 hour at 37 ℃ with 5 μM Cal520-AM (AAT Bioquest). Cells were then returned to conditioned media preserved prior to adding the calcium sensitive dye and allowed to equilibrate for >30 min at 37 ℃ and 5% CO_2_. Following equilibration, tissues were imaged under epifluorescence at 65 Hz while maintaining temperature and CO_2_. Time-lapse movies of calcium flux were analyzed with custom Matlab scripts as previously described^15^. Briefly, average fluorescent profiles over time were determined for each tissue and parameters such as beats per minute, peak-to-peak irregularity, flux rise time, flux decay time, and peak full width half max were calculated. Contraction correlation coefficient was determined by dividing the entire tissue into 16 regions of equal area and calculating the average Pearson’s correlation coefficient between the flux profiles of each of these regions.

### Immunofluorescence staining

Samples were fixed in 2% paraformaldehyde for 10 min at RT. Samples were then permeablized in PBS solution containing Triton X-100 (0.2% v/v), sucrose (10% w/v), and magnesium chloride (0.6% w/v) for 10 min and blocked in 1% (w/v) bovine serum albumin. Alternatively, to extract cytoplasmic vinculin, samples were simultaneously fixed and permeabilized in 2% paraformaldehyde in a buffer containing 1,4-piperazinediethanesulfonic acid (PIPES, 0.1 M), ethylene glycol-bis(2-aminoethylether)-N,N,N’,N’-tetraacetic acid (EGTA, 1 mM), magnesium sulfate (1 mM), poly(ethylene glycol) (4 % w/v), and triton X-100 (1% v/v) for 10 min at room temperature, prior to blocking in 1% (w/v) bovine serum albumin. Tissues were incubated with rabbit monoclonal anti-N-cadherin (1:500; Abcam Ab18203), rabbit monoclonal anti-connexin43 (1:1000; Millipore Sigma AB1728), mouse monoclonal anti-α-actinin (1:500; Abcam ab9465), mouse monoclonal anti-cardiac troponin T (1:500; ThermoFisher MA5-12960), mouse monoclonal anti-vinculin (1:1000; Millipore Sigma V9264), rabbit polyclonal anti-myosin light chain 2 (1:500; Proteintech 10906-1-AP), rabbit polyclonal anti-non-muscle myosin II-B (1:1000; Biolegend 909902), rabbit monoclonal anti-paxillin (1:500; Abcam ab246719), or mouse monoclonal anti-integrin β1D (1:1000; Abcam ab8991) antibodies for 1 hour at RT, followed by goat anti-mouse Alexa Fluor 647 (1:1000; Life Technologies A21236), goat anti-mouse Alex Fluor 546 (1:1000, Life Technologies A-11030), or goat anti-rabbit Alexa Fluor 647 secondary antibodies (1:1000; Life Technologies A21245) and DAPI for 1 hour at RT.

### Microscopy and image analysis

Fluorescent images were captured on a Zeiss LSM800 confocal microscope. Sarcomere alignment was quantified via custom Matlab scripts as previously described^15^. Briefly, images of titin-GFP reporter were thresholded and individual z-discs segmented. Z-discs were subsequently grouped with neighboring z-discs based on proximity and orientation to identify myofibrils within the image. The orientation of all identified myofibrils within a field of view was fit to a Gaussian distribution. Sarcomere alignment deviation was defined at the standard deviation of this distribution using circular/angular statistics. Myofibril density was calculated by determining the percent area of each tissue containing titin-rich myofibril structures. Vinculin morphology and colocalization analysis was also performed using custom Matlab scripts.

### Computational Modeling

The development of a tissue specific finite element model of fibroTUG tissues is described in detail by Jilberto et al. in an accompanying manuscript^49^. Briefly, the images of the DVS fibers and titin were processed using Matlab/Python scripts to quantify the specific fiber structure and a probabilistic characterization of the myofibril organization^69^. This information was projected into a 2D triangular finite element mesh. Using these quantities, and following a continuum mechanics approach, non-linear constitutive relationships for the fibers and cells were defined. The mechanical response of the tissue was taken to be the sum of these two components. We then adapted methods similar to those presented in Miller et al. (2021)^70^ to find the necessary active stress that the cells are exerting to generate the observed boundary tractions and displacement conditions. Using the experimental data for each of the permutations of interest (soft/stiff fibers with soft/stiff post), we generated more than a hundred *in-silico* tissues for each of them by combining image-derived fibrous matrix with probabilistic-generated myofibril fields and experimentally measured force responses specific for each condition. We then compile the results to obtain a distribution of active stress for each mechanical environment.

### Statistical analysis

Statistical significance was determined by t-tests and one-way or two-way analysis of variance (ANOVA) with post-hoc analysis (Tukey test), where appropriate, with significance indicated by p < 0.05. Studies were conducted a minimum of 3 times in each experiment. The data presented are representative data sets from one of these replicate studies. Specific sample size is indicated within corresponding figure legends and all data are presented as mean ± standard deviation.

## DATA AVAILABILITY

All data generated or analyzed during this study are available within the article and its supplementary information files or from the corresponding author upon reasonable request.

