## Supplemental Figures S1-S8 for "Matrix architecture and mechanics regulate myofibril organization, costamere assembly, and contractility of engineered myocardial microtissues"

Assistant Professor, Department of Biomedical Engineering, University of Michigan

2174 Lurie BME Building, 1101 Beal Avenue

Ann Arbor, MI 48109

*SUPPLEMENTAL FIGURES*

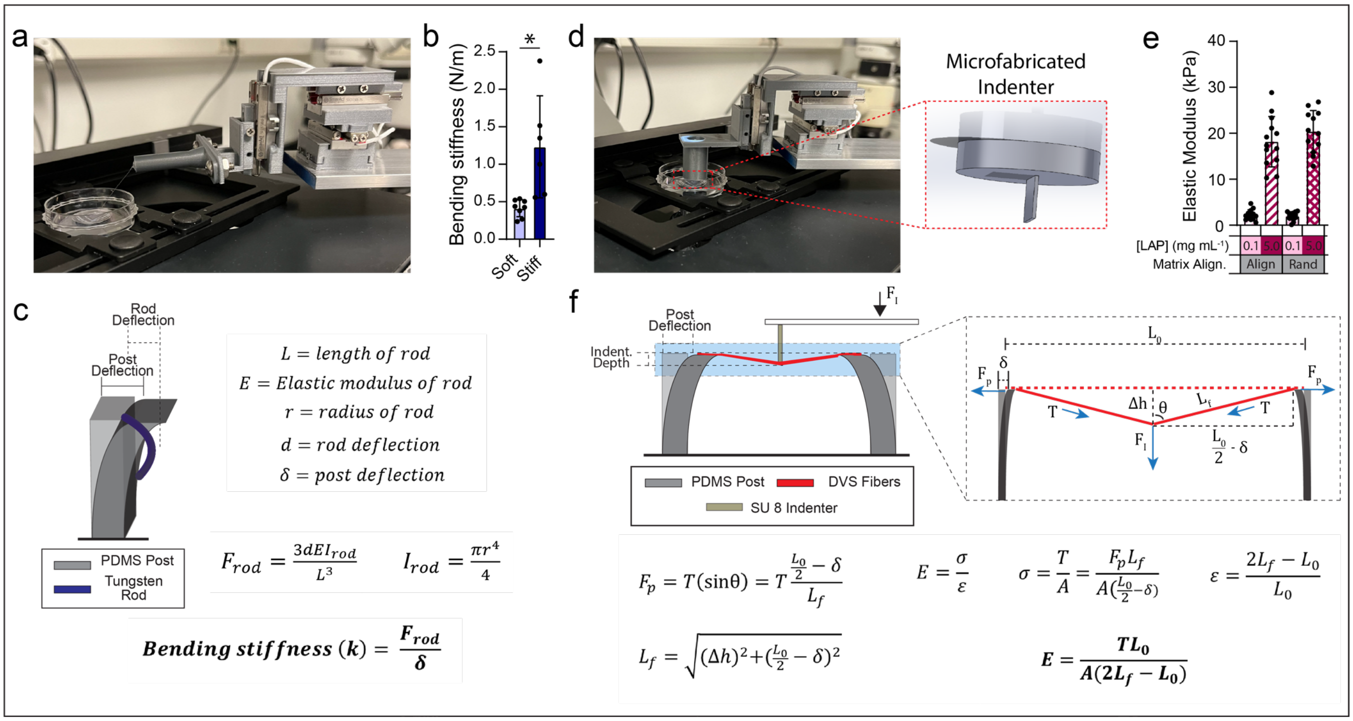


**Figure S1: Mechanical characterization of fibroTUG platform.** (a) Image of PDMS post mechanical characterization setup. (b) Measured bending stiffness of soft and stiff PDMS posts. (c) Schematic of post mechanical testing scheme and equations used to calculate post bending stiffness. (d) Image of fiber matrix mechanical characterization setup. (e) Measured elastic modulus of various fiber matrices. (f) Schematic of fiber matrix mechanical testing scheme and equations used to calculate matrix elastic modulus. All data presented as mean ± std; * $p < 0.05$.

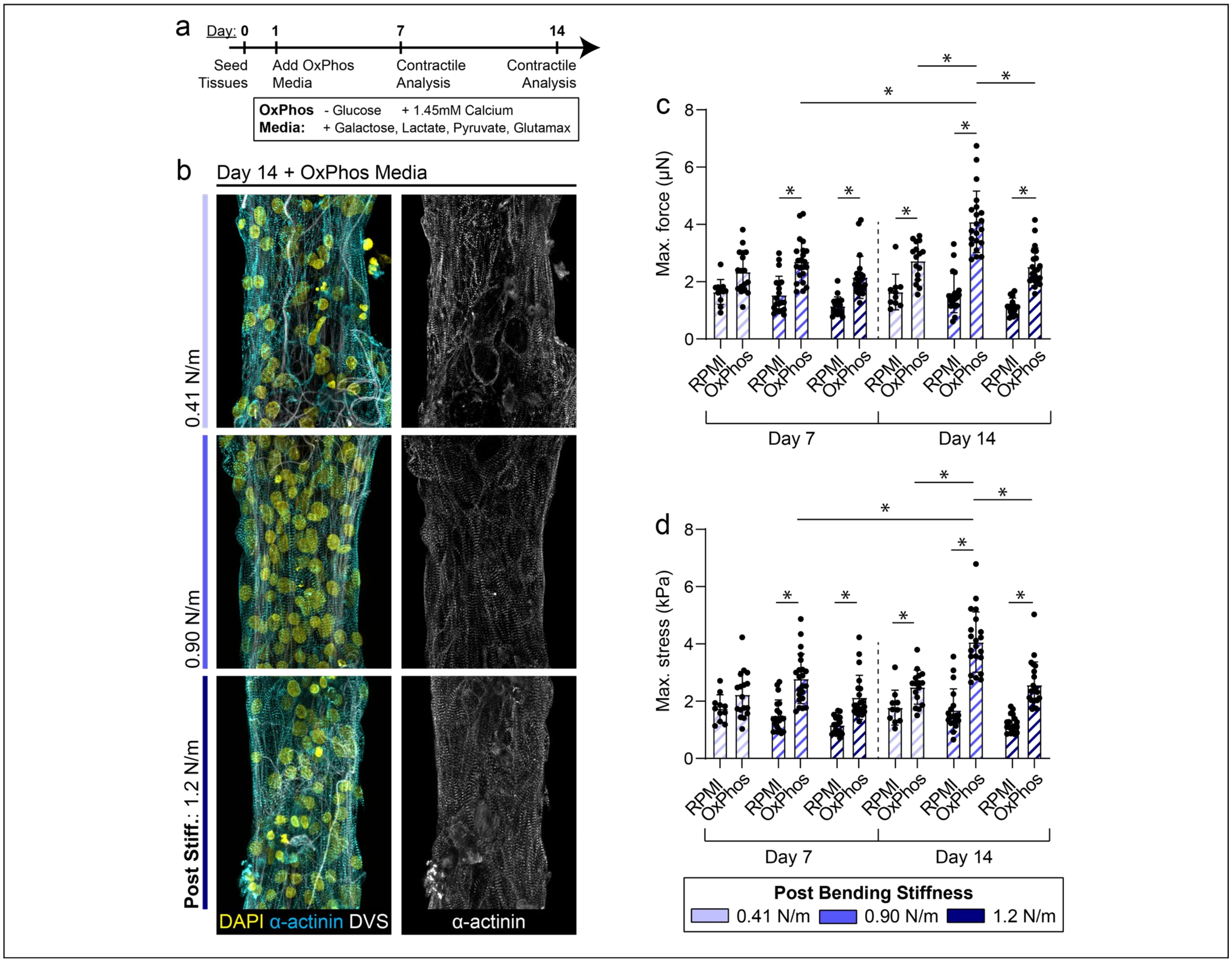


**Figure S2: fibroTUG platform supports long-term culture of iPSC-CMs with metabolic maturation media.** (a) Experimental timeline. (b) Confocal fluorescent images of fibroTUG tissues after 14 days in culture with OxPhos metabolic maturation media. (c) Max contractile force and (d) max contractile stress of tissues between various post stiffnesses in both RPMI B27 and OxPhos media. All data presented as mean ± std; * $p < 0.05$.

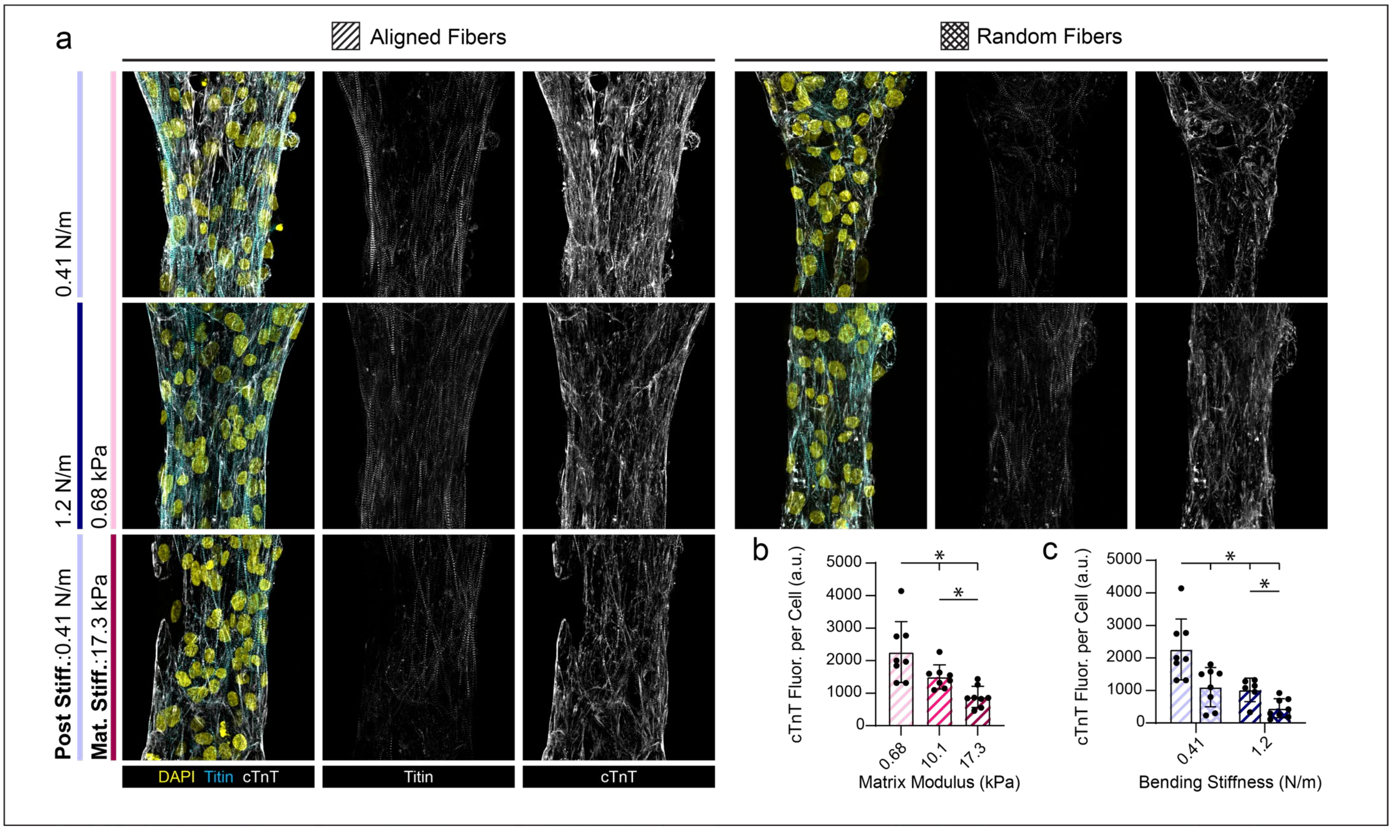


**Figure S3: Cardiac troponin expression in fibroTUG tissues.** (a) Confocal fluorescent images of fibroTUG tissues of varying mechanics immunostained for cTnT. Quantification of cTnT fluorescence per cell in tissues with (b) varied matrix stiffness and (c) bending stiffness. All data presented as mean ± std; * $p < 0.05$.

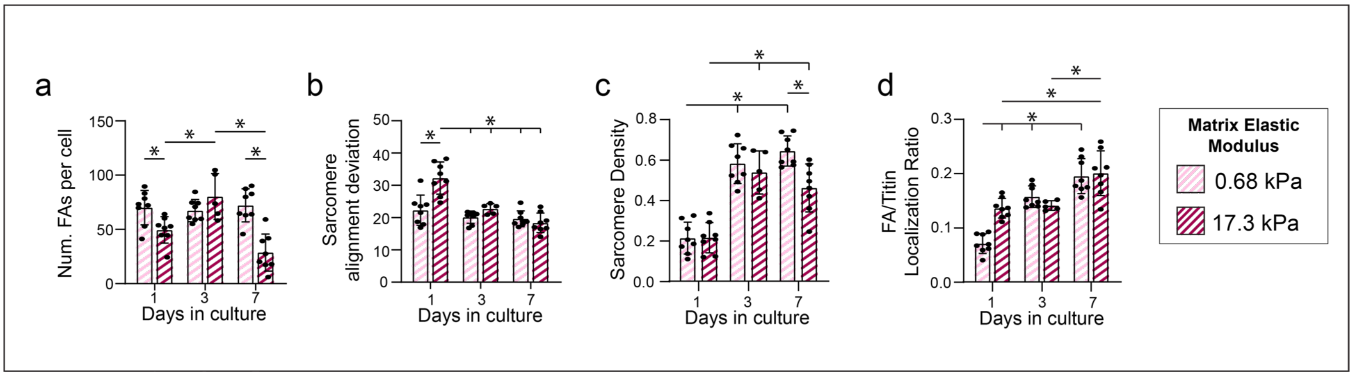


**Figure S4: Matrix mechanics influence costamere formation and sarcomere formation over time.** Quantification of number of (a) focal adhesions per cell, (b) sarcomere alignment, (c) sarcomere density, (d) and the fraction of vinculin that is colocalized with titin in tissues formed on aligned soft and stiff matrices suspended between 0.41 N/m posts fixed at 1-, 3-, and 7-days post seeding ($n \geq 5$). All data presented as mean ± std; * $p < 0.05$.

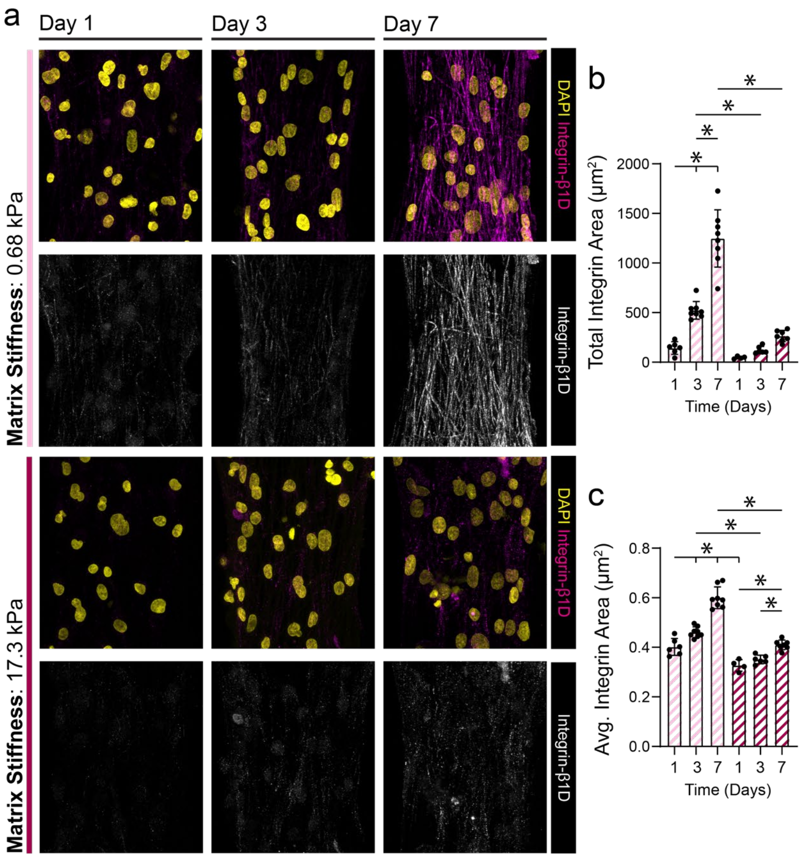


**Figure S5: Quantification of β1D integrin expression in fibroTUG tissues.** (a) Confocal fluorescent images of fibroTUG tissues fixed at day 1, 3 and 7 post seeding on either soft (0.68 kPa) or stiff (17.1 kPa) aligned fiber matrices (post stiffness was held constant at 0.41 N/m). Quantification of (b) total integrin area and (c) average integrin size ($n \geq 5$). All data presented as mean ± std; * $p < 0.05$.

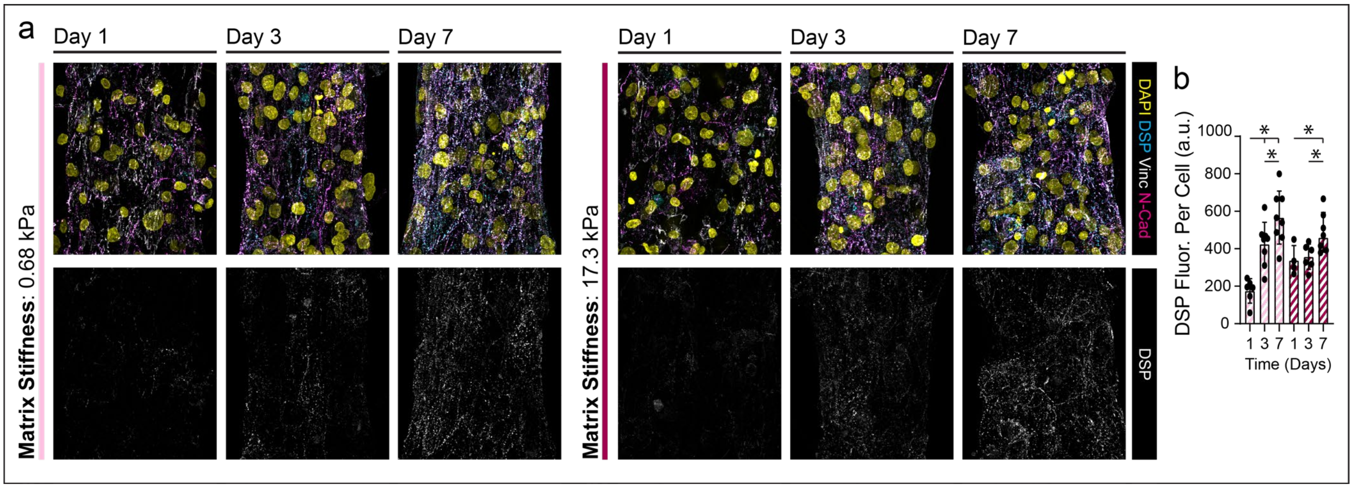


**Figure S6: Intercalated disc formation over time on soft and stiff fiber matrices.** (a) Confocal fluorescent images of fibroTUG tissues fixed at day 1, 3 and 7 post seeding on either soft (0.68 kPa) or stiff (17.1 kPa) aligned fiber matrices (post stiffness was held constant at 0.41 N/m). (b) Quantification of DSP fluorescence per cell ($n \geq 8$). All data presented as mean ± std; * $p < 0.05$.

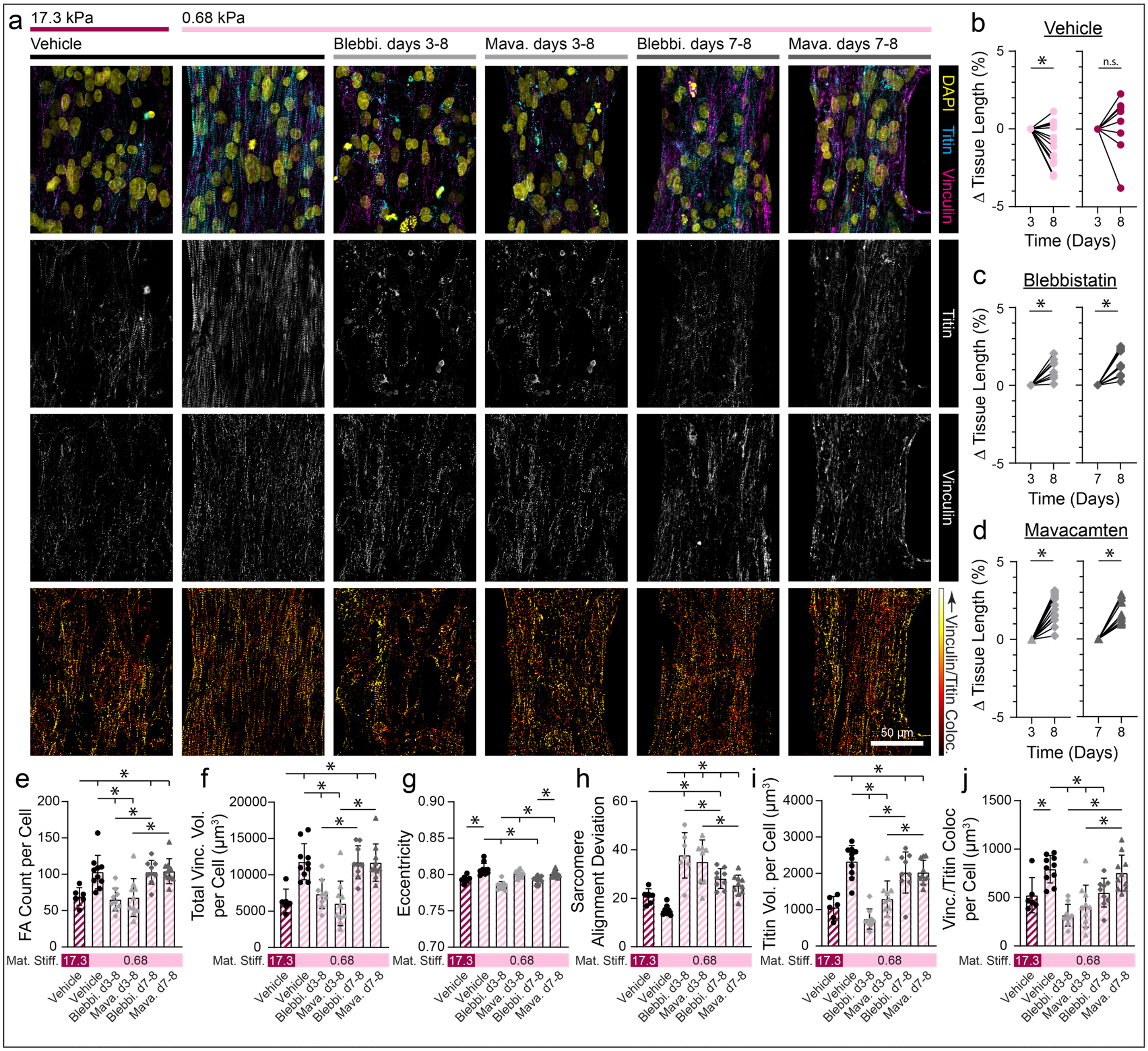


**Figure S7: Treatment with contractile inhibitors at day 3 and day 7.** (a) Confocal fluorescent images of fibroTUG tissues treated with blebbistatin (50uM) or mavacamten (500 nM) starting at either day 3 or day 7post seeding. (b) Diastolic tissue length on day 3 and 8 of tissues seeded on soft (0.68 kPa) and stiff (17.1 kPa) aligned fiber matrices (post stiffness was held constant at 0.41 N/m) without treatment with the contractile inhibitors. (c,d) Diastolic tissue length of tissues seeded on soft matrices on day 3 or day 7 before treatment with a contractile inhibitor, (c) blebbistatin or (d) mavacamten, and day 8 after 5 or 1 days of treatment,($n \geq 8$). (e) Focal adhesion count, (f) vinculin volume per cell, and (g) focal adhesion eccentricity were quantified from the fluorescent images of immunostained vinculin ($n \geq 6$). (h) Sarcomere alignment deviation and (i) titin volume per cell quantified from fluorescent images of titin-GFP reporter ($n \geq 6$). (j) Vinculin colocalization with titin per cell quantified from titin and vinculin images ($n \geq 6$). All data presented as mean ± std; * $p < 0.05$.

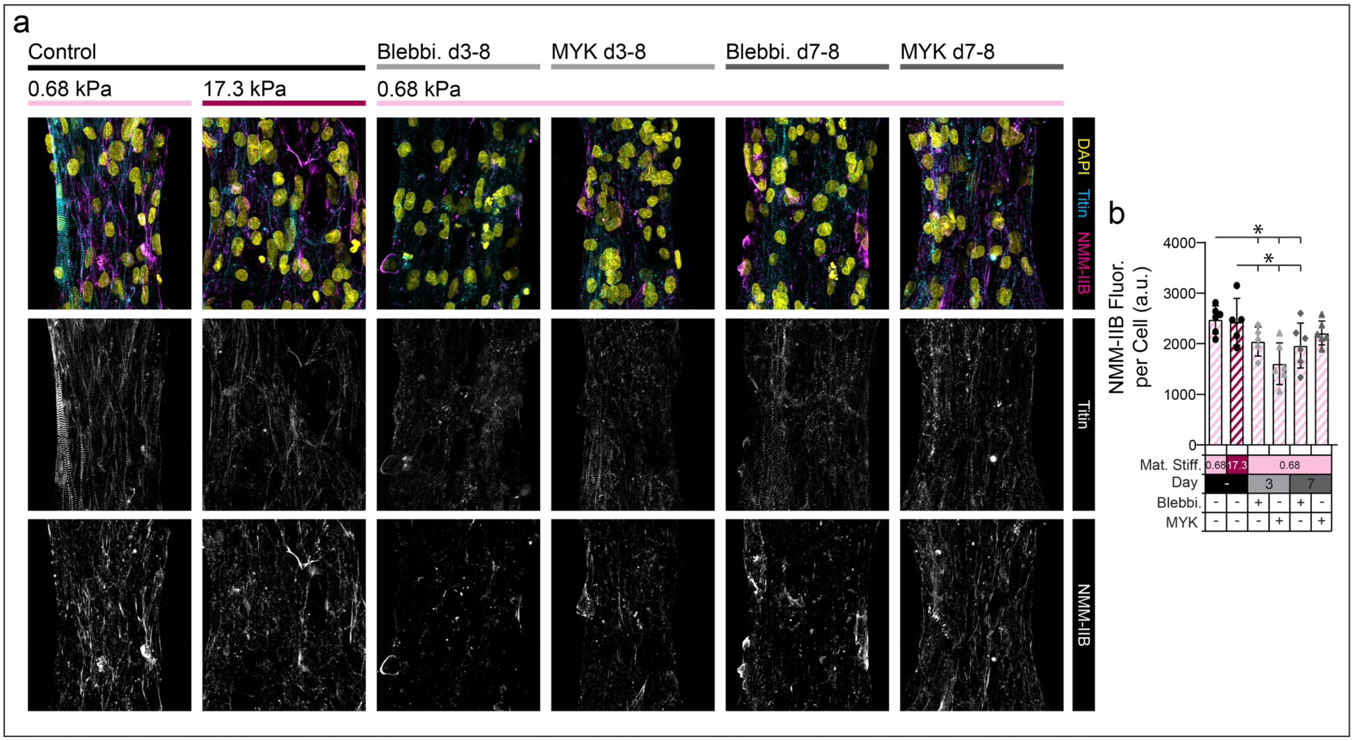


**Figure S8: Treatment with myosin inhibitors at day 3 decreases non-muscle myosin IIB (NMM-IIB) expression. (**a) Confocal fluorescent images of fibroTUG tissues treated with blebbistatin (50 µM) or mavacamten (500 nM). (b) Quantification of NMM-IIB expression per cell ($n \geq 6$). All data presented as mean ± std; * $p < 0.05$.

### SUPPLEMENTAL VIDEOS

**Supplemental Video S1:** Representative brightfield videos of contracting fibroTUG tissues formed on aligned fiber matrices of varying stiffness suspended between soft (0.41 N/m) posts.

**Supplemental Video S2:** Representative brightfield videos of contracting fibroTUG tissues formed on random fiber matrices of varying stiffness suspended between soft (0.41 N/m) posts.

**Supplemental Video S3:** Representative brightfield videos of contracting fibroTUG tissues formed on aligned, soft (0.68 kPa) fiber matrices suspended between posts of varying stiffness.

**Supplemental Video S4:** Representative brightfield videos of contracting fibroTUG tissues formed on random, soft (0.68 kPa) fiber matrices suspended between posts of varying stiffness.

**Supplemental Video S5:** Representative videos of calcium fluxes in fibroTUG tissues treated with Cal520-AM dye formed on aligned fiber matrices of varying stiffness suspended between soft (0.41 N/m) posts.

**Supplemental Video S6:** Representative videos of calcium fluxes in fibroTUG tissues treated with Cal520-AM dye formed on aligned, soft (0.68 kPa) fiber matrices suspended between posts of varying stiffness.

**Supplemental Video S7:** Representative videos of calcium fluxes in fibroTUG tissues treated with Cal520-AM dye formed on random, soft (0.68 kPa) fiber matrices suspended between posts of varying stiffness.

**Supplemental Video S8:** Representative brightfield videos of contracting fibroTUG tissues formed on aligned, soft (0.68 kPa) fiber matrices suspended between soft (0.41 N/m) posts before and after isoproterenol treatment.
